# HnRNP C binding to inverted *Alu* elements protects the transcriptome from pre-mRNA circularization

**DOI:** 10.1101/2025.08.04.666350

**Authors:** Alberto Marini, Consuelo Pitolli, Sabrina Ciccone, Marco Pieraccioli, Noémie Robil, Chiara Naro, Fernando Palluzzi, Manuela Giansanti, Gianpiero Tamburrini, Luciano Giacò, Pierre de la Grange, Francesca Nazio, Claudio Sette, Vittoria Pagliarini

**Affiliations:** Department of Neuroscience, Section of Human Anatomy, Università Cattolica del Sacro Cuore, 00168 Rome, Italy; GSTEP-Organoids Research Core Facility, IRCCS Fondazione Policlinico Universitario Agostino Gemelli, 00168 Rome, Italy; Genosplice, Paris, France; Bioinformatics Research Core Facility, Gemelli Science and Technology Park (GSTeP), IRCCS Fondazione Policlinico Universitario Agostino Gemelli, 00168 Rome, Italy; Innate Lymphoid Cells Unit, Bambino Gesù Children’s Hospital IRCCS, Rome, Italy; Pediatric Neurosurgery, IRCCS Fondazione Policlinico Universitario Agostino Gemelli, 00168 Rome, Italy; Department of Biology, University of Rome Tor Vergata, Rome, Italy

**Keywords:** RNA-binding proteins, inverted repeated elements, circular RNA, back-splicing

## Abstract

Back-splicing is a non-canonical splicing event that drives the biogenesis of circular RNAs (circRNAs). Although the molecular mechanisms underlying circRNA biogenesis have been partially elucidated, how this process is globally regulated in tumors, has not been fully investigated. Herein, we uncover a hnRNP C-dependent mechanism that represses a broad repertoire of circRNAs in Group 3 medulloblastoma (MB). HnRNP C binds *Alu* elements and prevents the circularization of pre-mRNA transcripts. Expression of hnRNP C modulates the balance between linear and circular splicing and guarantees efficient expression of genes that sustain the oncogenic phenotype of Group 3 MB cells. Remarkably, in the absence of hnRNP C, the introns flanking the circularizing exons generate cytoplasmic dsRNAs through base-pairing of inverted *Alu* elements and trigger an interferon-induced antiviral response. These findings unveil the role of hnRNP C as guardian of transcriptome integrity by repressing circRNA biogenesis. Lastly, targeting hnRNP C in Group 3 MB may trigger an inflammatory immune response, thereby boosting cancer surveillance.

## INTRODUCTION

Circular RNAs are covalently closed molecules generated by a back-splicing reaction in the precursor messenger RNAs (pre-mRNAs) (1). Back-splicing involves the splicing between a downstream 5’ splice site with an upstream 3’ splice site and, as canonical linear splicing, is mediated by the spliceosome machinery and regulated by similar sequence elements and RNA-binding proteins (RBPs) (1–3). The resulting circRNAs are generally more stable than their linear counterparts, and have been shown to affect many biological processes, ranging from transcription to translation of target mRNAs or protein functions in both nucleus and the cytoplasm (1–3). Notably, dysregulation of circRNA expression has been linked to many human diseases, particularly to human cancers (3, 4). Introns that flank circularizing exons are generally characterized by increased length and are enriched in inverted *Alu* repeat elements (1–3).These elements are essential to form a transient *Alu* pairing-mediated double-stranded RNA (dsRNA) between the involved introns, thus bringing into proximity back-splicing sites that are otherwise distant from each other and favoring the back-splicing event (1–3, 5). In addition to inverted *Alu* sequences, RBPs also play a fundamental regulatory role in circRNA biogenesis, by binding in proximity of back-splicing sites and modulating the efficiency of the reaction (6–9). Thus, it is conceivable that the specific repertoire of RBPs greatly influences the circRNAs that are expressed into each given cells. Alteration of this balance often occurs in disease states, such as cancer, where the expression of RBPs is often deregulated (10). In this context, RBPs have been shown to either promote the expression of oncogenic circRNAs or, alternatively, to inhibit circRNA biogenesis, thus ensuring the proper processing and expression of pro-tumoral linear transcripts that are essential for tumor cell survival. Interestingly, mounting evidence suggests that tumors are characterized by a global downregulation of circRNA expression with respect to healthy tissues (11, 12). Herein, we found that the RBP hnRNP C is a critical regulator of the balance of circRNA biogenesis in the context of medulloblastoma (MB), the most common malignant brain tumor of childhood (13, 14). Notably, MB arises in the cerebellum, the brain region with the highest expression levels of circRNAs (15). Among the different MB subgroups, Group 3 tumor displays the most aggressive phenotype and shorter survival (13, 14). Amplification and/or overexpression of the oncogenic transcription factor MYC drives an abnormal transcriptional program in Group 3 MB cells. Interestingly, MYC-amplified tumors are characterized by an increased transcriptional rate and accumulation of nascent pre-mRNAs in the nucleus, thus imposing a stress to the splicing machinery (16). As a consequence, MYC-amplified tumor cells are generally dependent on an efficient splicing machinery for their survival (16). To support the splicing machinery, MYC promotes the expression of a vast repertoire of splicing factors in many tumor cell types (17–20). Since linear splicing and back-splicing compete for the same pre-mRNA (21–23), it is conceivable that the global repression of circRNA biogenesis in tumor cells fuels the increased output of canonical mRNAs in high demanding tumor cells, such as those harboring amplification of MYC. Thus, we reasoned that the regulatory mechanism(s) controlling the extent of circRNA biogenesis might be particularly exacerbated in MYC-amplified Group 3 MB. In line with this hypothesis, RNA sequencing (RNA-seq) analysis of samples from high-risk Group 3 MB patients revealed a strong downregulation of circRNAs with respect to healthy individuals. Motif enrichment analyses and functional studies demonstrated that hnRNP C exerts a widespread inhibitory effect on circRNA biogenesis in Group 3 MB cells, while concomitantly supporting the expression of cognate linear transcripts. Mechanistically, we show that hnRNP C preferentially binds proximal intronic *Alu* elements that flank the circularizing exons, thus inhibiting the occurrence of the back-splicing event. In the absence of hnRNP C, paired intronic dsRNAs are stabilized and, after the back-splicing reaction, eventually accumulate into the cytoplasm and trigger the activation of the innate immune response, contributing to the reduced fitness of MB cells. Together, our findings uncover a functional role of hnRNP C as a general negative regulator of circRNA biogenesis and suggest the existence of a hnRNP C-dependent safeguard mechanism to repress aberrant circularization of pre-mRNAs in MYC-amplified tumors.

## RESULTS

### CircRNA expression is extensively downregulated in Group 3 MB patients

Large transcriptomic analyses in different types of cancers, including MB, highlighted a significant downregulation of circRNAs compared to normal tissues (11, 12). This observation suggests a functional correlation between repression of circRNA biogenesis and tumorigenesis. To identify and quantify circRNAs expressed in high-risk Group 3 MB, we searched for back splicing junctions (BSJs) by querying RNA-seq data downloaded from the International Cancer Genome Consortium (ICGC) Controlled Data Portal (dataset EGAD00001004958). We compared 16 Group 3 MBs with 4 fetal (FC) and 5 adult cerebella (AC), thus including both undifferentiated and differentiated healthy tissues (Supplementary Table S1). Computational analyses carried out with CircExplorer2 and CIRI, two widely used tools for circRNA detection (24, 25), identified 12343 BSJs expressed in at least one of the experimental groups, with 3444 high confidence events supported by both tools (Fig. 1A; Supplementary Table S2). Among these common BSJs, more than 90% are predicted to yield exonic circRNAs (Supplementary Fig. S1A). While a fraction of these BSJs (n=923; 26.8%) were detected in at least two experimental groups, the majority (n=2521; 73.1%) were specifically detected only in one of them (Fig. 1B; Supplementary Table S2). Most of these unique BSJs were detected either in the AC (n=1461) or in the MB (n=880) samples. Differential expression analysis of BSJs in these two groups showed that circRNAs are mostly downregulated in our cohort of Group 3 MB patients compared to AC (Fig. 1C; Supplementary Table S3), while no differences were observed with respect to FC (Supplementary Fig. S1B; Supplementary Table S4).

**Figure 1.**
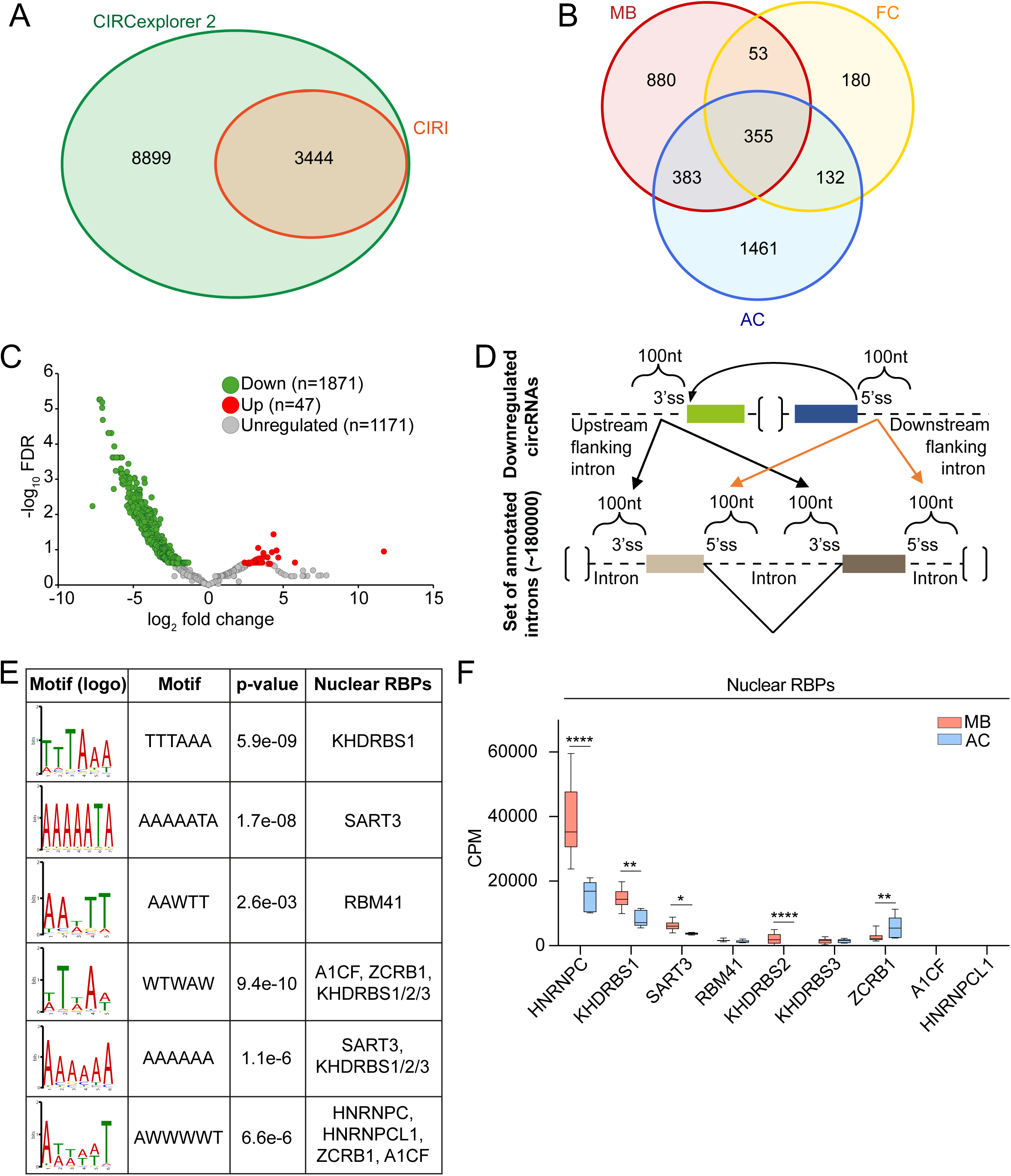
CircRNA expression is extensively downregulated in Group 3 MB patients. (**A**) Identification of circRNAs querying EGAD00001004958 RNA-seq dataset. Venn diagram showing the intersection between CircExplorer2 and CIRI (n=3444) computational analyses.. (**B**) Venn diagram depicting the number of circRNAs identified in healthy cerebella (AC, n=5; FC, n=4) and Group 3 MB (MB, n=16). (**C**) Differential expression analysis of circRNAs detected in Group 3 MB and adult cerebellum (AC). circRNAs differentially expressed (FDR<0,25) were highlighted in red and green. (**D**) Scheme depicting the strategy for the motif enrichment analysis using STREME-TomTom (MEME suite, https://meme-suite.org/meme/) in circRNAs flanking introns. (**E**) Motifs and corresponding RBPs predicted in motif enrichment analysis in panel D. (**F**) mRNA expression levels (CPM, count per million reads) in EGAD00001004958 RNA-seq data of nuclear RBPs whose binding sites were enriched (Benjamini–Hochberg adjusted p-values; *p < 0.05, **p < 0.01, ****p < 0.0001).

To investigate whether suppression of circRNA biogenesis in Group 3 MB is an active regulatory mechanism, we explored the *trans*-acting factors that are potentially involved in back-splicing modulation. To this end, we searched for RNA sequence motifs enriched in the introns flanking the back-spliced exons of AC unique circRNAs (n=1461; Fig. 1B). Using the MEME suite tool (26), we compared the sequence of regions encompassing 100 nt upstream of the 3’ splice sites or downstream of the 5’ splice sites, involved in the circularization of AC-unique exons, with the corresponding intronic regions not involved in back-splicing events (∼180000 intron sequences, UCSC genome browser- https://genome.ucsc.edu; Fig. 1D; Supplementary Table S5). STREME analysis (27) identified putative consensus motifs enriched near the unique AC BSJs, while the TomTom Motif Comparison tool (28, 29) predicted the RBPs that potentially bind these motifs (Fig. 1E). Taking into consideration the top 3 most significantly enriched motifs upstream and downstream the BSJs, we identified nine nuclear RBPs (KHDRBS1/Sam68, SART3, RBM41, A1CF, ZCRB1, KHDRBS2/SLM1, KHDRBS3, HNRNPC, and HNRNPCL1) as potential regulators of AC unique exons back-splicing (Fig. 1E). Next, to identify MB-specific RBPs involved in the biogenesis of these circRNAs, we focused on RBPs that were significantly upregulated in our cohort of Group 3 MB patients compared to healthy individuals (Fig. 1F and Supplementary Fig. S1C; Supplementary Table S6). Based on these inclusion criteria, we selected HNRNPC, KHDRBS1/Sam68, SART3, and KHDRBS2/SLM1 for subsequent analysis. Notably, all these RBPs showed the highest protein expression in MB compared to other pediatric brain tumors analyzed (Supplementary Fig. S1D-G), further suggesting their involvement in this cancer type. Gene expression analysis in public MB datasets confirmed the upregulation of hnRNP C, SART3, Sam68 and SLM1 in Group 3 MB (n=233) compared to normal cerebellum (n=291). By contrast, SLM1 was preferentially upregulated in Group 3 and 4 MB respect to SHH and WNT MB (Supplementary Fig. S1H-K). Interestingly, hnRNP C and SART3 are the only RBP specifically upregulated in Group 3 MB patients also with respect to the other MB subgroups [Group 4, Sonic Hedgehog (SHH) and Wingless (WNT)] (Supplementary Fig. S1G,H).

### HnRNP C represses the expression of circRNAs in Group 3 MB cells

Previous studies reported a role for hnRNP C and Sam68 in the regulation of circRNA biogenesis (7, 30), whereas no information is available for SART3 and SLM1. To validate the functional role of the selected RBPs in circRNA biogenesis in Group 3 MB cells, we knocked down their expression in D341-Med (hereafter D341) and HD-MB03 cells. Silencing efficiency was evaluated by western blotting for hnRNP C, SART3 and Sam68 (Fig. 2A; Supplementary Fig. S2B) whereas, due to cross-reactions of the SLM1 antibody with the highly homologous Sam68 (Fig. 2A and Supplementary Fig. S2B), SLM1 expression was evaluated by qPCR (Supplementary Fig. S2A,C). To test the impact of these RBPs on circRNA biogenesis, we selected five MB expressed circRNAs (circMTDH, circCACNA2D1, circFIP1L1, circHNRNPM, circRBM28) that harbor the binding sites for all 4 RBPs in the intronic regions flanking the BSJ (Fig. 2B-F and Supplementary Fig. S2D-H; Supplementary Table S7). Strikingly, we observed that knockdown of hnRNP C was sufficient to significantly induce the expression of all five circRNAs in Group 3 MB cells. Notably, increased circRNA expression was also accompanied by the concomitant reduction of the cognate linear transcript in D341 cells (Fig. 2B-E), indicating that hnRNP C may modulate the competition between back- and linear splicing in these target genes.

**Figure 2.**
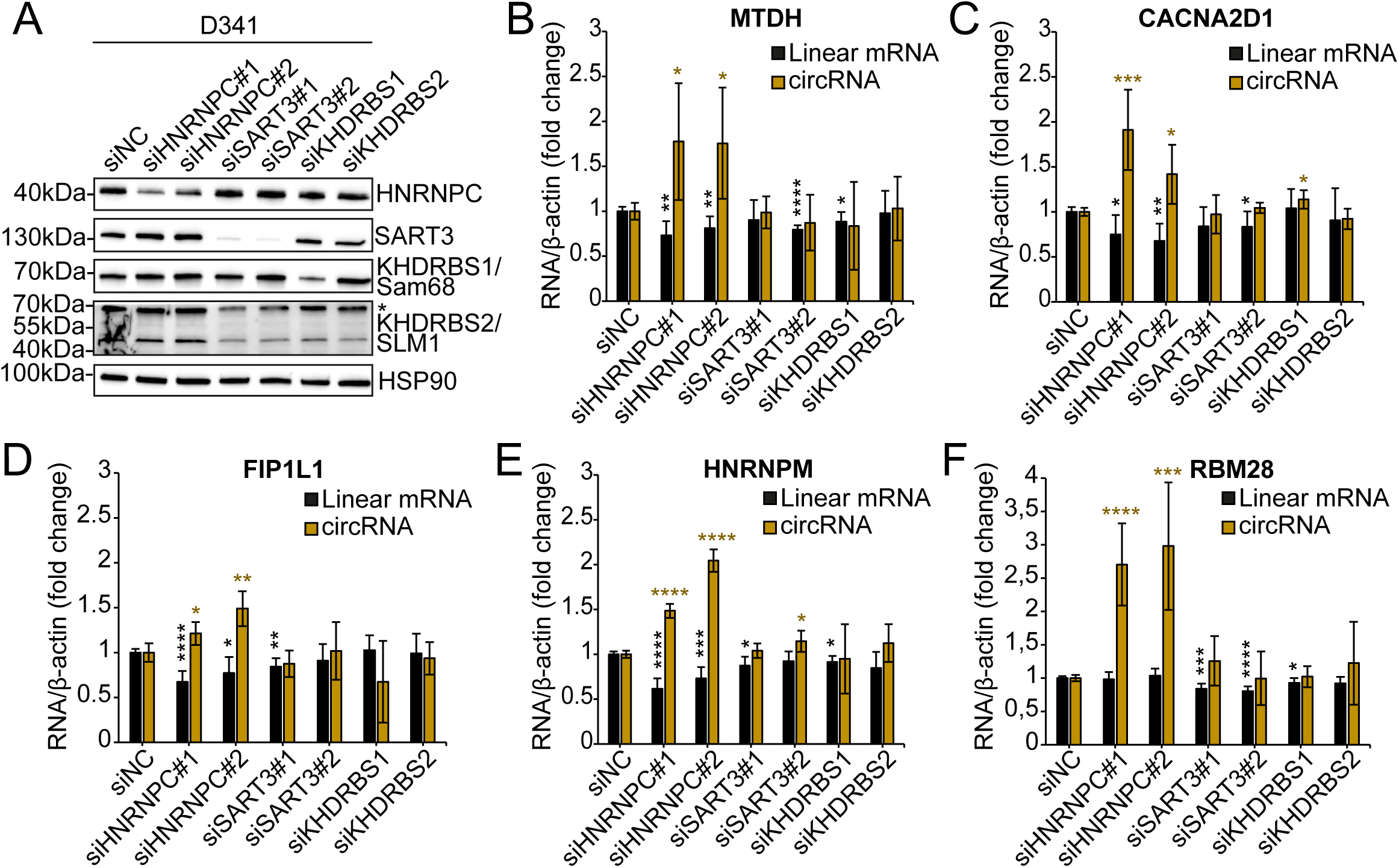
HnRNP C represses the expression of circRNAs in Group 3 MB cells. (**A**) Western blot analysis evaluating the expression of the selected RBPs upon their silencing in D341 MB cells. Asterisk (*) indicates a not-specific signal for KHDRBS1/Sam68 in KHDRBS2/SLM1 western blot. (**B-F**) qPCRs detecting parental linear mRNAs and circRNAs having enrichment for HNRNPC, SART3, and KHDRBS1/2 binding sites in the flanking introns. #1 and #2 indicate two different siRNAs. siNC: siRNA negative control (n=3; mean ± SD; Student’s t-test vs siNC; *p < 0.05, **p < 0.01, ***p < 0.001, ****p < 0.0001).

By contrast, knockdown of the other three RBPs caused only mild or no effects on the expression of these circRNAs (Fig. 2B-F; Supplementary Fig. S2D-H). Importantly, the same effect of hnRNP C knockdown was observed by using two different siRNAs in both cell lines (Fig. 2A and Supplementary Fig. S2B), except for circCACNA2D1 that was only significantly upregulated in D341 cells (Fig. 2C). Together, these results suggest that hnRNP C plays a functional role in the negative regulation of circRNA biogenesis in Group 3 MB cells.

To further explore the functional role of hnRNP C as general repressor of circRNA biogenesis, we carried out RNA-seq analyses in D341 cells. Treatment of the RNA samples with RNaseR (Supplementary Fig. S3A,B) was performed to enrich for circRNAs while removing linear RNAs (31). Principal component analysis (PCA) and unsupervised clustering of RNAseR-treated cells confirmed the segregation of control and hnRNP C-depleted samples (Supplementary Fig. S3C). Analysis of RNaseR-treated samples by the CIRI software identified a total of 19291 BSJs in D341 cells (Supplementary Table S8). In agreement with previous findings (5), introns flanking the exons undergoing back-splicing in D341 cells are longer and contain more inverted and repeated *Alu* (IR*Alu*) elements than introns not surrounding circularization events (Supplementary Fig. S3D,E; Supplementary Tables S9,S10). Of note, introns flanking BSJs were also enriched for stretches of ≥9 uridine residues (Supplementary Fig. S3F; Supplementary Table S11), which are strong binding motifs for hnRNP C (32). To directly determine the role of hnRNP C on circRNA biogenesis, we then focused on 3479 high confidence back-splicing events from 2050 genes which showed at least two read counts in at least two replicates of either control (siNC) or hnRNP C-depleted (siHNRNPC) D341 cells (Supplementary Table S8). The vast majority (94.7%) of these back-splicing events involved exons (Fig. 3A; Supplementary Table S8). Notably, knockdown of hnRNP C induced a robust increase in the number of unique BSJs (∼4-fold, Fig. 3B) and in circRNA-producing genes (3.5-fold, Fig. 3C). Furthermore, comparison of back-splicing and linear splicing in transcripts expressed in all samples (n=1685; Supplementary Table S8) showed a highly significant increase of circRNA:linear RNA ratio in hnRNP C-depleted cells (Fig. 3D), suggesting that hnRNP C modulate the balance between linear and circular splicing in these cells. Moreover, transcriptome analyses of parallel samples not treated with RNAseR indicated that depletion of hnRNPC has a significantly stronger impact on the modulation of circRNAs (17% of expressed genes) than on canonical splicing (6%) and gene expression (5%) (Fig. 3E, Supplementary Fig. S3G, Supplementary Tables S12,S13).

**Figure 3.**
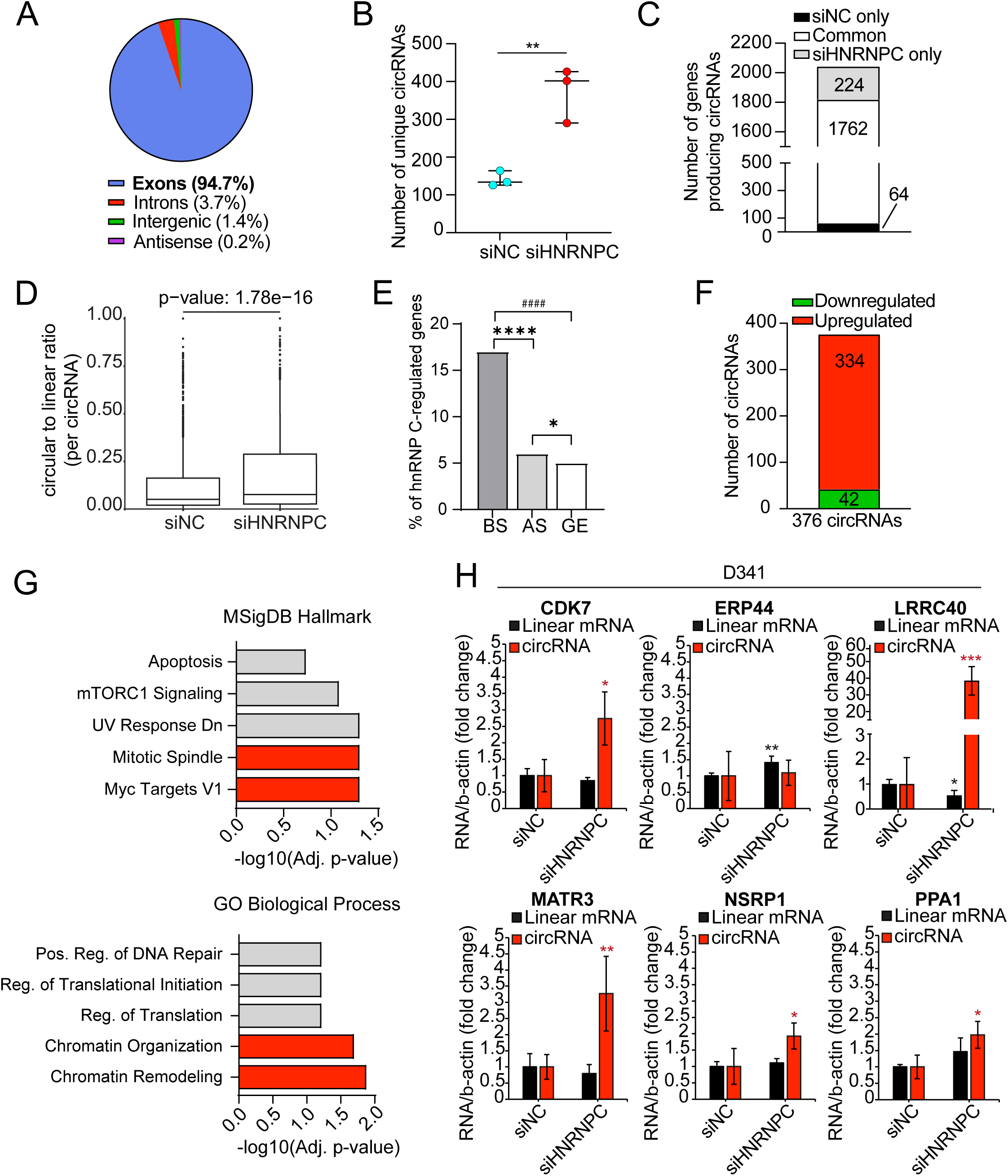
HnRNP C is a general repressor of circRNA biogenesis in Group 3 MB cells. (**A**) Percentage of circRNAs generated from exons, introns, intergenic regions, and antisense transcripts in siNC and siHNRNPC samples. (**B**) Number of unique circRNAs detected in siNC and siHNRNPC treated D341 cells. Median with interquartile range is shown (Student’s t-test; **p < 0.01). (**C**) Number of circRNA-producing genes in siNC and siHNRNPC samples. (**D**) Circular to linear read-count ratio per circRNA in control and HNRNPC depleted cells. p-value is shown (Student’s t-test). (**E**) Percentage of hnRNP C-regulated genes in terms of back-splicing (BS; n=352), alternative splicing (AS; n=648) and gene expression (GE; n=572). Fisher’s exact test; *p=0.03, ****p=8.17e^-51^, ^####^p=6.5e^- 60^. (**F**) Number of up-regulated and down-regulated circRNAs upon HNRNPC depletion. (**G**) Gene set enrichment analysis of the genes generating up-regulated circRNAs was performed using Enrichr (https://maayanlab.cloud/Enrichr/). (**H**) Validation of 6 up- regulated exonic circRNAs contained in the list of the top 8 most up-regulated circRNAs in HNRNPC depleted cells (n=4; mean ± SD; Student’s t-test vs siNC; *p < 0.05, **p < 0.01, ***p < 0.001).

Our analysis indicated that depletion of hnRNP C affects 376 circRNAs in D341 cells, with ∼90% of them being upregulated (Fig. 3F; Supplementary Tables S8,S14). Notably, analysis of the genes generating the up-regulated circRNAs (n=311) by the Molecular Signatures Database (MSigDB) uncovered a significant enrichment in MYC target and mitotic spindle genes (Fig. 3G, upper panel). Furthermore, Gene Ontology analyses highlighted a significant enrichment in genes associated with chromatin remodeling and positive regulation of DNA repair (Fig. 3G lower panel), two biological processes involved in tumor evolution and resistance to chemotherapy. Together, these findings suggest that hnRNP C is required to repress excessive circularization of transcripts that are crucial for Group 3 MB cell viability and proliferation. Analysis by RT-qPCR validated the increased expression of 5/6 and 6/6 of the upregulated circRNAs in, respectively, D341 and HD- MB03 cells (Fig. 3H; Supplementary Fig. S3H; Supplementary Table S14).

### Introns flanking circularizing exons are enriched in strong binding sites for hnRNP C in close proximity of *Alu* elements

Analyses of intron sequence features indicated that those flanking hnRNP C-repressed BSJs are shorter (Fig. 4A; Supplementary Tables S8,S15) and characterized by increased frequency of IR*Alu* elements and hnRNP C binding sites with respect to introns surrounding unregulated back-splicing events (Fig. 4B,C; Supplementary Tables S8,S15,S16). Moreover, the hnRNP C binding sites are found at the beginning of intronic *Alu* elements, regardless of regulation, but they are closer to the regulated BSJs than the non-regulated ones (Fig. 4D,E; Supplementary Tables S17,S18). Thus, hnRNP C preferentially represses back-splicing events mediated by introns with IR*Alu* close to the splice sites.

**Figure 4.**
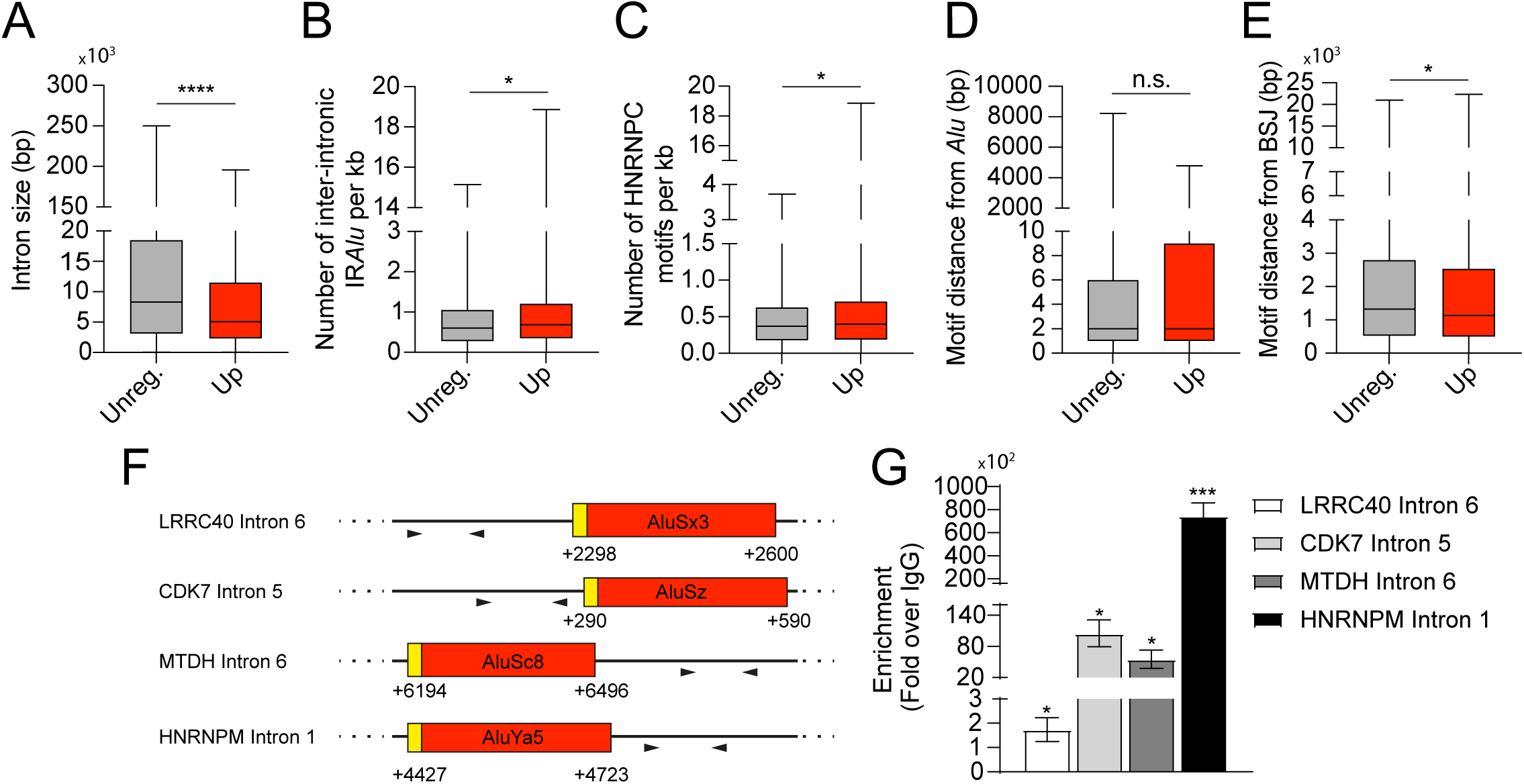
Introns flanking circularizing exons are enriched for hnRNP C strong binding sites in proximity of *Alu* elements. (**A-E**) Features of the flanking introns sequences belonging to the up-regulated circRNAs in HNRNPC depleted cells compared to flanking introns of unregulated circRNAs. Student’s t-test; *p < 0.05, ****p < 0.0001, n.s. stands for not significant. (**F**) Schematic representation of hnRNP C binding sites analyzed by CLIP-qPCR within the indicated transcripts. Yellow boxes indicate predicted hnRNP C binding motifs overlapping with *Alu* elements (shown in red). Arrowheads denote the positions of primer pairs used in CLIP-qPCR to quantify pre-mRNA region bound by hnRNP C. Nucleotide positions are given relative to the intron start site, where the first nucleotide of the intron is designated as +1. (**G**) CLIP assays performed in D341 cells using anti-hnRNP C antibody or IgGs as a negative control. Data are represented as fold change enrichment over IgGs (n=3; mean ± SEM; Student’s t-test vs IgG; *p < 0.05, ***p < 0.001).

To test whether hnRNP C binds introns involved in the regulated back-splicing events, we performed UV crosslink immunoprecipitation (CLIP) experiments. Strikingly, strong binding of hnRNP C was observed in close proximity of *Alu* elements within the introns flanking the regulated BSJs, where its binding sites were predicted to be located (Fig. 4F,G). These observations support a key role of hnRNP C as general repressor of circRNA biogenesis through the direct binding in proximity of *Alu* elements in Group 3 MB cells.

### Binding of hnRNP C is necessary to inhibit back-splicing events

We hypothesized that hnRNP C binding at the beginning of the *Alu* elements may interfere with IR*Alu* pairing between the introns flanking circularizing exons, thus repressing circRNA biogenesis. To test this hypothesis, we took advantage of a *Survival Motor Neuron 2* (*SMN2*) circRNA-producing minigene previously validated in our laboratory (7). First, we observed that hnRNP C depletion caused the upregulation of the endogenous circSMN9-6 in D341 cells (Fig. 5A). Next, we employed the *SMN2* minigene encompassing the coding region of the *SMN2* gene from exon 5 to the region downstream of exon 8, which comprises the cryptic exon 9 involved in the back-splicing event with exon 6 (7). An internal portion of intron 6 was deleted to reduce the size of the minigene, while a six-nucleotide TAG was inserted in exon 6 to allow discrimination of the recombinant RNA products from endogenous transcripts (Fig. 5B). *SMN2* intron 5, which represents the upstream intron involved in the back-splicing event, has all the sequence features highlighted by our bioinformatics analysis (Fig. 4A-E). First, it is a relatively small intron (1310 bp) comprising two *Alu* elements (AluSq; +364-671 bp and FLAM_C; +1051-1188 bp) and 2 strong binding sites for hnRNP C located at nucleotide +374 (binding site 1 (BS1); stretch of 13 uridine residues) and +879 (binding site 2 (BS2); stretch of 12 uridine residues). Both hnRNP C binding sites are located at the beginning of the *Alu* elements. Furthermore, BS2 is relatively close (420 bp) to the acceptor back-splicing site (Fig. 5B). Transfection of the minigene in HEK293T cells resulted in the production of the expected circSMN9-6 (Fig. 5C). Interestingly, hnRNP C depletion strongly increased the expression of circSMN9-6 (Fig. 5C), indicating that this minigene is suitable to study the hnRNP C-dependent regulation of circRNA biogenesis. To test whether binding of hnRNP C near the *Alu* elements is required to repress circRNA biogenesis, we mutagenized the minigene to disrupt the hnRNP C binding. To this aim, we replaced both binding sites for hnRNP C in intron 5, alone or in combination, by substituting the thymidine stretch with adenines (Fig. 5D and Supplementary Fig. S4). Strikingly, mutation of either hnRNP C binding site increased circSMN9-6, with BS1 showing a stronger effect. Moreover, combined disruption of both binding sites completely rescued circRNA biogenesis to the same levels of hnRNP C silencing (Fig. 5E). These results suggest that hnRNP C binding in proximity of *Alu* elements is necessary and sufficient to repress the SMN circRNA biogenesis.

**Figure 5.**
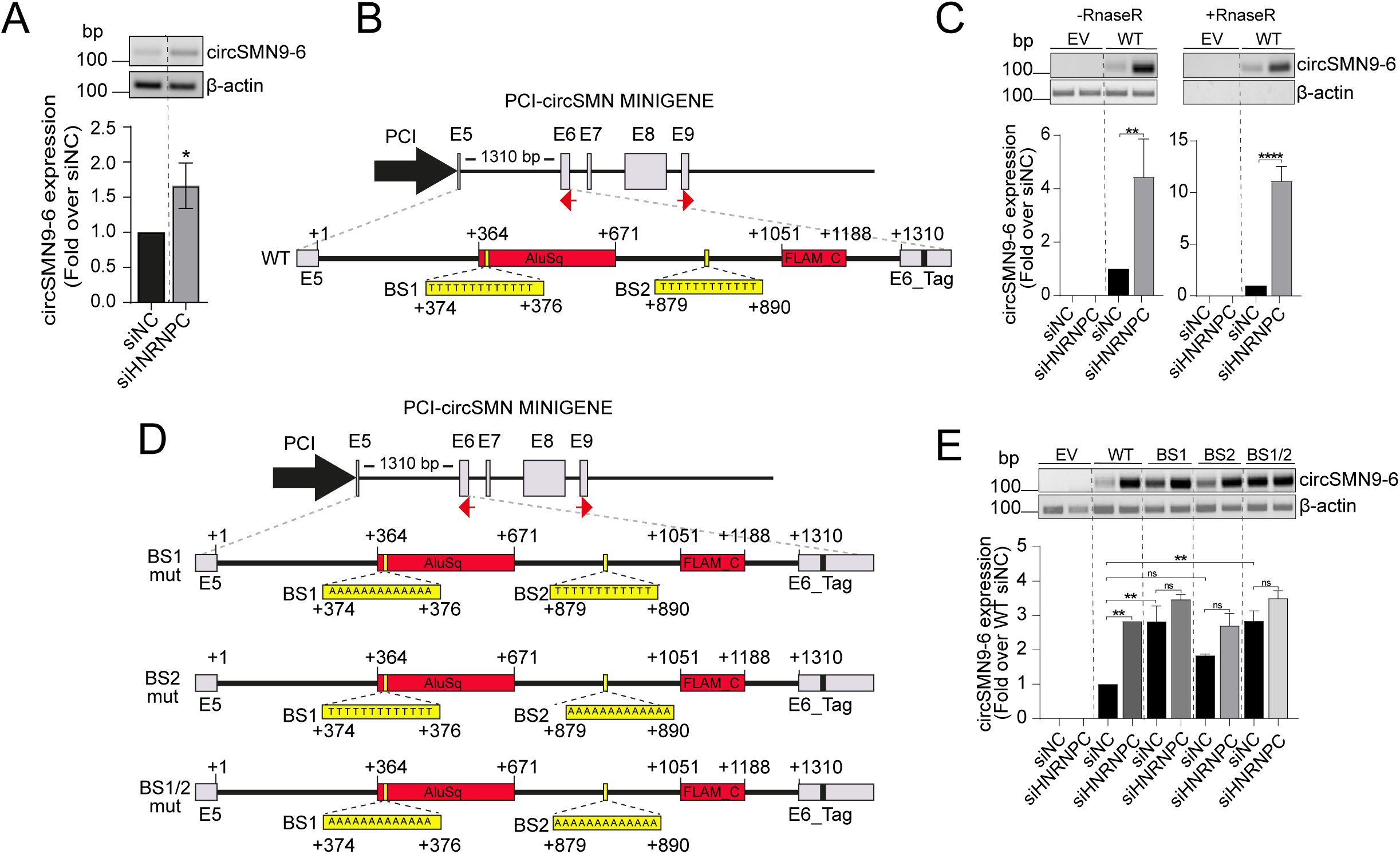
Binding of hnRNP C is necessary to inhibit back-splicing events. (**A**) Representative RT-PCR image (upper panel) and densitometric analysis (lower panel) showing endogenous circSMN9-6 expression in D341 cells transfected with siNC or siHNRNPC. Actin was used as a loading control (n=3; mean ± SD; Student’s t-test; p ≤ 0.05). (**B**) Schematic representation of the wild-type circSMN9-6 minigene (WT). Grey boxes indicate exons, red boxes represent *Alu* elements, black lines denote introns, and red arrows indicate divergent primers used for the detection of minigene-derived circSMN9-6. The nucleotide sequence highlights HNRNPC binding sites within SMN2 intron 5. (**C**) Representative RT-PCR image (upper panel) and densitometric analysis (lower panel) showing minigene-derived circSMN9-6 expression in HEK293T cells transfected with either empty vector or the circSMN9-6 WT minigene, with or without RNase R treatment. Actin was used to normalize RNA input (n=3; mean ± SD; One-way Anova; **p < 0.01, ****p < 0.0001). (**D**) Schematic representation of mutant circSMN9-6 minigenes. BS1, BS2, and BS1/BS2 indicate mutations in HNRNPC binding site 1, binding site 2, and both sites, respectively. (**E**) Representative RT-PCR image (upper panel) and densitometric analysis (lower panel) showing circSMN9-6 expression derived from mutant minigenes in HEK293T cells transfected with BS1, BS2, or BS1/BS2 constructs, following transfection with siNC or siHNRNPC. Actin was used as a loading control (n=3; mean ± SD; Student’s t-test; **p < 0.01). ns stands for not significant.

### HnRNP C represses the accumulation of intron-derived dsRNAs and spurious activation of innate immune response in Group 3 MB cells

The main mechanism underlying the regulation of circRNA biogenesis is the pairing of intronic IR*Alu* elements (5). We reasoned that these structures, if not properly controlled, may generate stable dsRNA adducts that could accumulate in the cell. To test this hypothesis, we carried out immunofluorescence assays with the J2 antibody, which specifically recognizes dsRNAs, in hnRNP C-depleted HD-MB03 cells, which grow in adherence and are more suitable for immunofluorescence analyses. Quantitative analysis of fluorescence intensity indicated that depletion of hnRNP C significantly increases the accumulation of cytoplasmic dsRNAs (Fig. 6A,B). The effect of hnRNP C knockdown on the accumulation of dsRNAs was similar to that elicited by DHX9 knockdown, a DNA/RNA helicase previously shown to globally inhibiting circRNA biogenesis by resolving the pairing of IR*Alu*-rich intronic regions and to repress cytosolic dsRNA accumulation (33).

**Figure 6.**
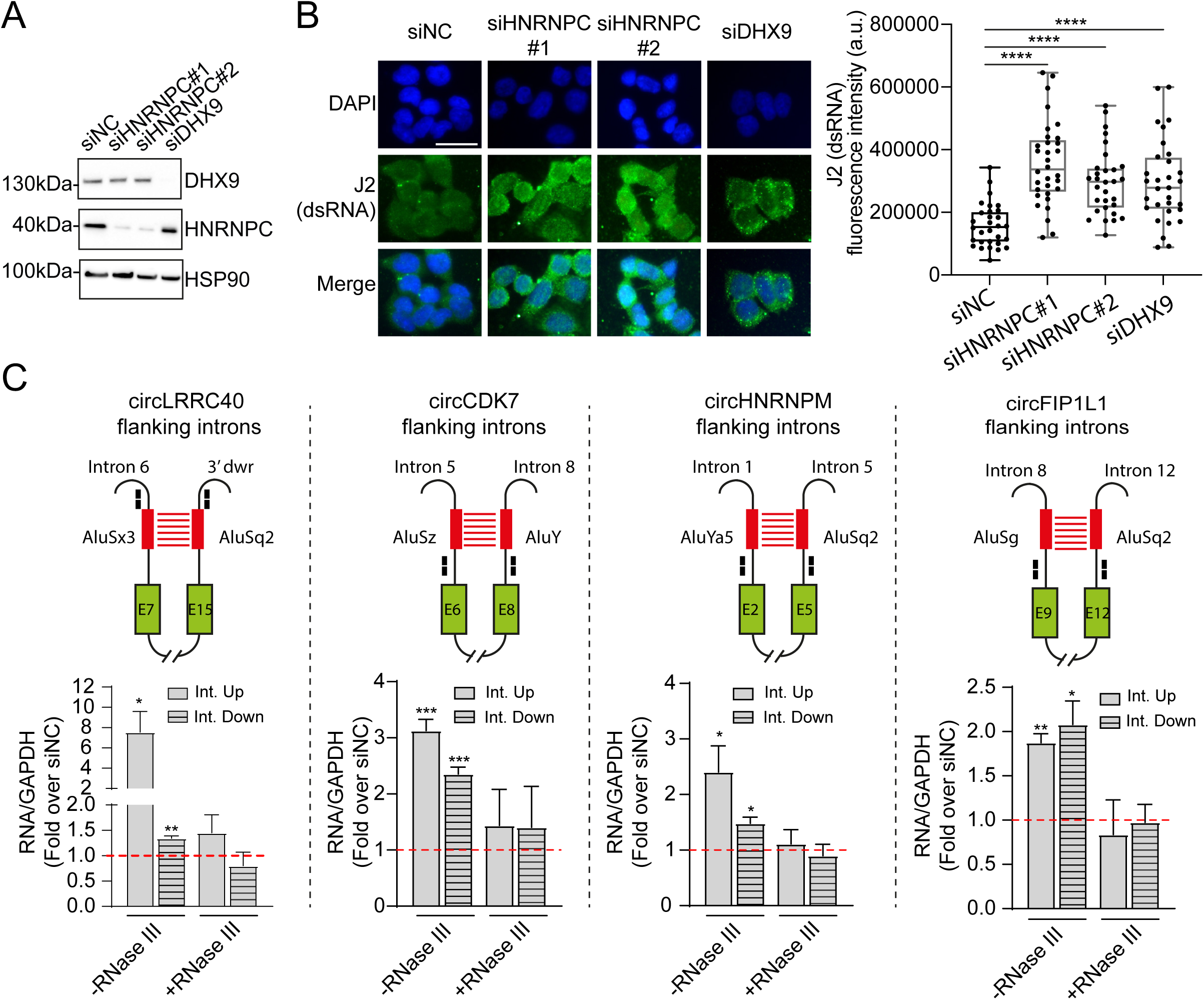
HnRNP C represses the accumulation of intron-derived dsRNAs in Group 3 MB cells. (**A**) Western blot analysis of HD-MB03 MB cells transfected with the indicated siRNAs. (**B**) Immunofluorescence analysis of dsRNA in HD-MB03 cells treated with the indicated siRNAs as in (A). J2 antibody was used for dsRNA detection. Scale bar: 20μm. Right panel shows quantification of cytoplasmic dsRNA signal (n=3; Student’s t-test vs siNC; ****p < 0.0001). (**C**) qPCR analysis of the flanking introns of the hnRNP C-regulated BSJs in the cytoplasmic fractions of hnRNP C-depleted HD-MB03 cells. Data are represented as fold change over siNC-treated cells set to 1, as indicated by the dashed red line. GAPDH was used as a loading control (n=3/4; mean ± SEM; Student’s t-test vs siNC; *p < 0.05, **p < 0.01, ***p < 0.001). Int. Up and Int. Down stand for upstream and downstream introns, respectively. The upper panel shows a schematic representation of the introns flanking the hnRNP C-regulated BSJs. Green boxes represent exons, black lines represent introns. Red boxes indicate *Alu* elements involved in the back-splicing event. Black boxes show the positions of primer pairs used for qPCR analysis.

To test whether intron flanking the hnRNP C-regulated BSJs contributed to such dsRNA accumulation, we performed sub-cellular fractionation assays in control and hnRNP C-depleted Group 3 MB cells (Supplementary Fig. S5A). RT-qPCR analysis by using primers in proximity to *Alu* sequences in introns flanking 4 arbitrarily selected BSJs revealed the accumulation of these introns in the cytoplasmic fractions of hnRNP C-depleted cells (Fig. 6C). Importantly, treatment of the extracts with RNase III, an endonuclease that degrade dsRNAs while preserving single strand RNAs (34), significantly reduced the accumulation of these intronic sequences in the cytoplasm (Fig. 6C), suggesting that these BJS-flanking introns are organized as cytoplasmic dsRNAs. In support of this notion, analyses of two introns not predicted to be involved in circRNA biogenesis did not accumulate in the cytoplasm in hnRNP C-depleted HD-MB03 cells (Supplementary Fig. S5B). These data uncover a new source of cytoplasmic dsRNAs as by-products of excessive circRNA biogenesis and a key role on hnRNP C in preventing accumulation of these nucleic acids adducts.

The accumulation of cytoplasmic dsRNAs in hnRNP C-depleted HD-MB03 cells positively correlated with the up-regulation of interferon-stimulated genes (ISGs) (i.e., TNF, IFNB1, CXCL10, IFI27 and IFI44; Fig. 7A). Unexpectedly, DHX9 depletion increased the expression of only two ISGs among those tested (CXCL10 and IFI44), suggesting that, at least in HD-MB03 cells, hnRNP C depletion triggers a stronger interferon response than DHX9 (Fig. 7A). IFNB1, IFI27, and IFI44 up-regulation was observed also in D341 cell (Supplementary Fig. S5C,D). Accordingly, hnRNP C depletion impaired cell viability in D341 and HD-MB03 cell lines (Fig. 7B,C) and reduced MB spheroid formation in D341 cells (Fig. 7D,E).

**Figure 7.**
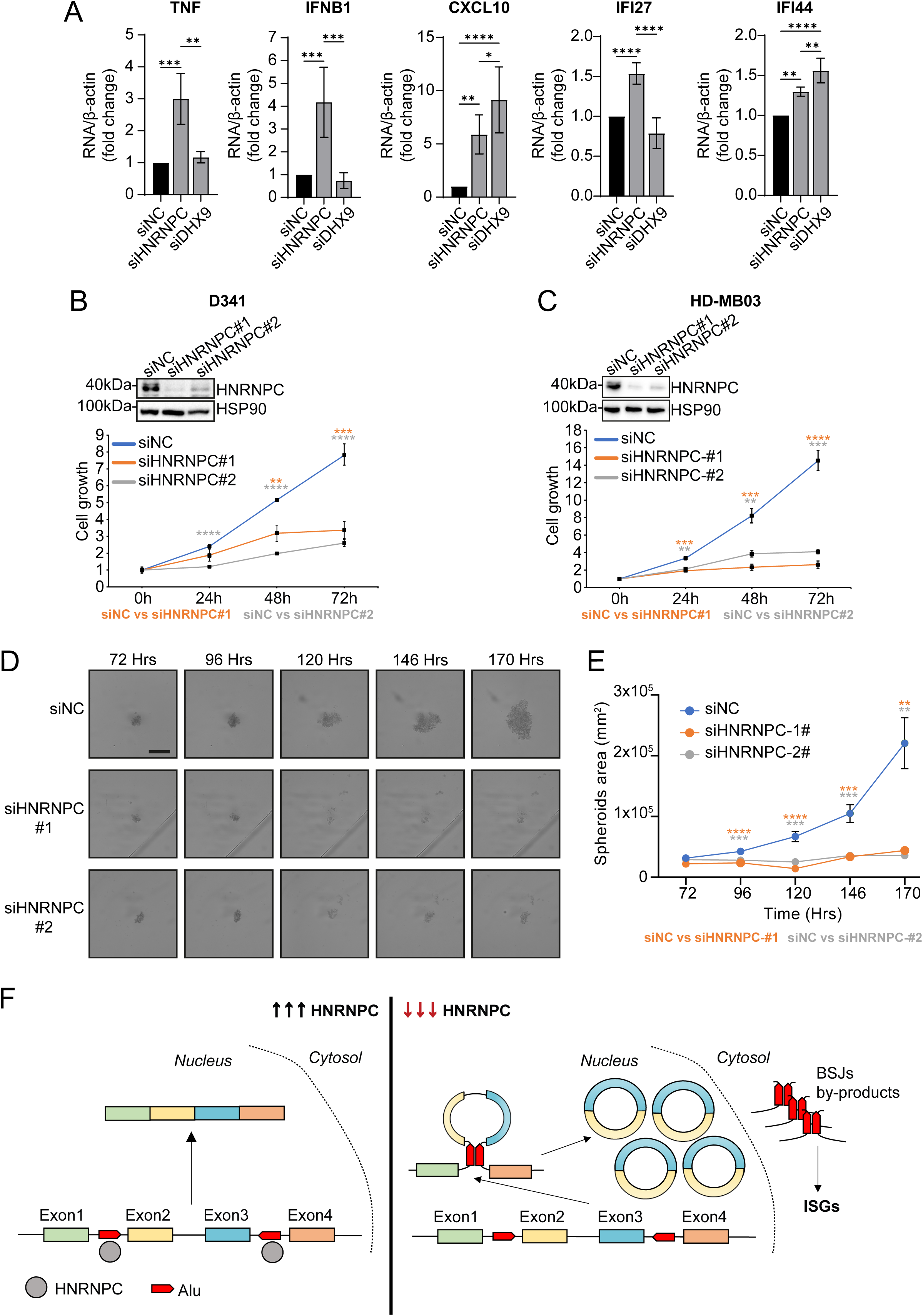
HnRNP C represses the spurious activation of innate immune response in Group 3 MB cells. (**A**) qPCR analysis of immune-related genes in HD-MB03 cells treated with the indicated siRNAs (n=3; mean ± SD; One-way ANOVA, *p < 0.05, **p < 0.01, ***p < 0.001, ****p < 0.0001). (**B**) Cell growth analysis in D341 cells transfected with the indicated siRNAs (n=3; Student’s t-test vs siNC; *p < 0.05, **p < 0.01, ***p < 0.001, ****p < 0.0001). (**C**) Cell growth analysis in HD-MB03 cells transfected with the indicated siRNAs (n=3; mean ± SD; Student’s t-test vs siNC; *p < 0.05, **p < 0.01, ***p < 0.001, ****p < 0.0001). (**D**) Microscope images of D341 spheroids at the indicated time points after seeding. Scale bar: 500µm. (**E**) Quantification of spheroids area at the indicated time points (n=12; mean ± SEM; Student’s t-test vs siNC; *p < 0.05, **p < 0.01, ***p < 0.001, ****p < 0.0001). (**F**) Schematic model showing the suppressive role of HNRNPC on circRNA expression in Group 3 MB, preventing the accumulation of cytoplasmic dsRNAs.

Together, these results highlight a novel role for hnRNP C as general repressor of IR*Alu* pairing and pre-mRNA circularization, which contributes to the safeguard of the human transcriptome integrity and to control inappropriate activation of the innate immune response.

## DISCUSSION

CircRNA expression is strongly inhibited in cancer cells respect to healthy tissues (11, 12). CircRNAs are generally more stable than linear transcripts and tend to accumulate in long-lived and post-mitotic cells, like neurons. At the same time, however, circRNA biogenesis has lower efficiency than production of canonical mRNAs through linear splicing (1). Thus, the lower content of circRNAs in cancer cells might represent the consequence of their higher proliferation rate and dilution of these less efficiently produced transcripts. However, mounting evidence also suggest that inhibition of circRNA biogenesis is important to sustain proliferation and viability of tumor cells (4). Herein, we identify hnRNP C as an important negative regulator of circRNA biogenesis in Group 3 MB. First, we observed a strong downregulation of circRNA expression in samples from Group 3 MB patients compared to adult healthy cerebellum. Furthermore, circRNA downregulation in Group 3 MB samples correlated with the higher expression levels of hnRNP C. By contrast, fetal healthy cerebellum, which express comparable levels of hnRNP C (Supplementary Fig. S1C), also expressed low levels of circRNAs, supporting a role for this RBP in the regulation of circRNA biogenesis in the healthy and pathological cerebellum. Moreover, we found that depletion of hnRNP C expression in Group 3 MB cells strongly increases the expression of hundreds of circRNAs, supporting the role of this RBP as a general repressor of circRNA biogenesis. These observations suggest that proliferative nervous tissues, such as fetal cerebellar precursor cells and MB cells, are more dependent on linear splicing for the production of large amounts of mRNAs and require high expression of hnRNP C to modulate the balance in favor of canonical splicing. This scenario might be particularly relevant for MYC-amplified Group 3 MB, due to the MYC-dependent transcriptional overload and the consequent higher dependence on an efficient splicing machinery for guaranteeing cell viability (16). Notably, another recurrent aberration in Group 3 MB is the amplification of the oncogenic transcription factor OTX2 (13) which was shown to induce the expression of hnRNP C (35). OTX2 physically interacts with hnRNP C and other members of the large assembly of splicing regulators (LASR) complex to drive an AS program that is essential to maintenance of stemness potential and progression of Group 3 MB (35). This experimental evidence further supports the strong dependence of MYC- and/or OTX2-amplified Group 3 MB from canonical splicing. In this sense, hnRNP C might function both to execute a cancer-related splicing program and, concomitantly, to prevent the usage of splice sites in non-canonical events, such as back-splicing.

Our unbiased search for RNA binding motifs enriched in the introns flanking the regulated BSJs has identified hnRNP C, KHDRBS1/Sam68, SART3 and KHDRBS2/SLM1 as potential regulator of circRNA biogenesis in Group 3 MB cells. All four RBPs were highly expressed in Group 3 MB patients at transcript and protein level. However, only hnRNP C depletion clearly impacted on circRNA biogenesis. Previous data indicated the possible implication of hnRNP C as regulator of pre-mRNA circularization. In particular, hnRNP C was shown to regulate the expression of three hypoxia-regulated circRNAs (circCDYL2, circRARS, and circSMARCA5) in HeLa cells, with two of them being repressed and one promoted by this RBP (30). However, to our knowledge, our study shows for the first time that hnRNP C acts as a general regulator of circRNA biogenesis with possible functional implications in MB context. Interestingly, for some pre-mRNA targets we show that hnRNP C depletion cause a concomitant increase in circularization and decrease in the linear counterpart, pointing to a key role of this RBP in the well-known competition between the canonical linear splicing and back-splicing (21–23). We posit that hnRNP C might function by repressing the formation of IR*Alu*-mediated secondary structures in the pre-mRNA that are necessary to promote non-colinear splicing, thus ensuring linear processing of transcripts. This hypothesis is supported by our bioinformatics analyses showing that the circularization index - the relative abundance of circRNA versus spliced linear mRNA – is significantly higher in hnRNP C-depleted cells, highlighting their higher proficiency to promote back-splicing of nascent pre-mRNAs. This role of hnRNP C as general repressor of circRNA biogenesis might be particularly relevant for Group 3 MB, which is frequently driven by oncogenic transcriptional factors (MYC, OTX2) that increase the transcriptional rate of the cell and may favor the prolonged life of intron-retaining transcripts, due to increased competition between introns for the splicing machinery. Importantly, the role of hnRNP C as repressor of IR*Alu*-mediated back-splicing is in line with its previously proposed role as repressor of cryptic intronic splice sites and exonization of transposable elements (36). Thus, hnRNP C may play a general role as guardian of the integrity of the human transcriptome by preventing aberrant exon inclusion and/or excessive circularization of pre-mRNAs comprising high density of transposable elements. Moreover, since genes affected by pre-mRNA circularization are enriched in terms of high relevance for Group 3 MB, such as MYC targets, mitotic spindle, chromatin remodelling and DNA repair (37), high expression of hnRNP C is likely required to favor linear splicing of transcripts that support oncogenic features of Group 3 MB cells.

Our bioinformatics analyses of RNAseR-treated Group 3 MB cells showed that the number of circRNAs and circRNA-generating genes was strongly increased in hnRNP C depleted cells. The impact of hnRNP C depletion on circRNA expression (17% of total) was significantly broader than on canonical splicing (6% of total) or gene expression (5% of total). Furthermore, ∼90% of these circRNAs were upregulated in the absence of hnRNP C. The introns flanking the regulated BSJs are significantly smaller and, more importantly, enriched for inter-intronic IR*Alu* and hnRNP C binding sites. Given that hnRNP C preferentially binds *Alu* elements (36), the higher number of hnRNP C-binding motifs in introns flanking BSJs might be due to the higher concentration of inter-intronic IR*Alu*. However, the hnRNP C-binding motifs are closer to the upregulated BSJs than to the unregulated BSJs. Thus, it is likely that hnRNP C represses the circularization of transcripts that, due to their intrinsic sequence features, tend to form IR*Alu*-mediated dsRNAs near a BSJ. Structurally, although hnRNP C contains only one RNA recognition motif (RRM), it oligomerizes into tetramers to form a complete functional unit for RNA binding (32). In line with this notion, ‘strong’ binding motifs for hnRNP C comprise stretches of ≥9 uridines (32). Coherently, we found such ’strong’ hnRNP C binding motifs at the beginning of *Alu* elements in both unregulated and regulated BSJs, which probably coincides with the terminal poly-A tail of the *Alu* element at the time of its insertion into the genome in opposite orientation. Previous experimental evidence showed the preferential binding of hnRNP C to antisense *Alu* elements (36). Furthermore, our *in vitro* mutagenesis experiments showed that mutation of the U-stretch in the predicted binding site for hnRNP C strongly increased the circularization of the circSMN2(ex9-6) and mimicked the effect of hnRNP C depletion. These results suggest that the direct binding of hnRNP C to U-stretches in proximity of IR*Alu* elements is required to repress the circularization of target transcripts. However, it is also conceivable that monomeric hnRNP C binds with lower affinity to shorter U-stretches along the entire length of *Alu* elements, thus contributing to further repress their pairing.

Our results also show that introns flanking hnRNP C-regulated BSJs accumulate into the cytoplasm as dsRNAs, thus activating the innate immune response. Since treatment with RNAse III completely abrogated their cytoplasmic accumulation, these RNA sequences from introns flanking the hnRNP C-regulated BSJs are likely structured in dsRNAs. Thus, we hypothesize that dsRNAs generated by BJS-flanking introns are byproducts of the increased circRNA biogenesis, which leak out into the cytoplasm and activate an innate immune response that hampers viability and proliferation of Group 3 MB-depleted cells. In this scenario, high expression of hnRNP C is required to maintain circRNA biogenesis under control and to limit the accumulation of dsRNAs generated by IR*Alu* pairing. This mechanism may be particularly relevant for tumors of tissues that accumulate large amounts of circRNAs, like the cerebellum. Thus, although a role for hnRNP C as repressor of innate immunity was already proposed in other cancer types (38, 39), herein we demonstrate for the first time that this dsRNA-induced antiviral response can be generated by pairing of intronic RNA sequences involved in back-splicing reactions. In support of this notion, our bioinformatics analyses did not show an enrichment in intron retention events into linear transcripts in hnRNP C-depleted cells that could justify the accumulation of cytoplasmic dsRNAs observed upon hnRNP C depletion (Supplementary Fig. S3G).

In conclusion, our study uncovers a key role of hnRNP C in maintenance of the transcriptome integrity by binding IR*Alu* elements in nascent RNAs and limiting their pairing and circRNA biogenesis. Thus, if on one hand hnRNP C prevents the exonization of *Alu* elements into the coding transcriptome (36), it also contributes to this process by repressing the circularization of the *Alu*-containing pre-mRNAs. In both cases, the lack of hnRNP C increases cytoplasmic accumulation of dsRNAs and activates a viral mimicry response. It remains to be investigated how these RNA adducts are transported from the nucleus into the cytoplasm, whether and how they are stabilized, and whether their specific structure (opened at both ends) requires a different repertoire of innate immunity sensors. Further studies will also be required to elucidate the functional role of hnRNP C- regulated circRNAs on the phenotype(s) of cerebellar cells and how their global reduction affects Group 3 MB cells. Moreover, as RBPs are emerging as therapeutic targets for diseases involving genomic abnormalities like cancer (40), inhibiting hnRNP C or enhancing the back-splicing-deriving dsRNAs could represent innovative methods to activate an immune response against MB.

## METHODS

### Human cell lines

D341-Med (D341), and HD-MB03 cells were cultured according to the recommended conditions (ATCC). D341 cells were cultured in Minimal Essential Medium (MEM, Gibco) supplemented with sodium pyruvate 1mM (Gibco), MEM Non-Essential Amino Acids Solution 1X (Gibco), 20% fetal bovine serum (FBS, Gibco), 100U/ml penicillin and 10μg/ml streptomycin (Euroclone). HD-MB03 cells (obtained from Deutsche Sammlung von Mikroorganismen und Zellkulturen DSMZ, Germany) were maintained in RPMI (Gibco) with 10% FBS (Gibco), 100U/ml penicillin and 100μg/ml streptomycin (Euroclone). HEK293T cells were maintained in Dulbecco-Minimal Essential Medium (DMEM, Sigma-Aldrich) supplemented with MEM Non-Essential Amino Acids Solution 1X (Gibco), 10% FBS (Gibco), 100U/ml penicillin and 100μg/ml streptomycin (Euroclone). All cell lines were cultured at 37°C in a humidified atmosphere with 5% CO_2_ and tested for mycoplasma contamination by PCR every 3 months.

### Cell transfections

Cells were transfected with 50nM of specific siRNAs using Lipofectamine RNAiMax Transfection Reagent (Cat. No. 13778150, Invitrogen) according to manufacturer’s instructions. The following siRNAs were used in this study: HNRNPC (siRNA#1, HSS179304; siRNA#2, HSS179305) (Stealth siRNAs ThermoFisher), SART3 (siRNA#1, SASI_Hs01_00115659; siRNA#2, SASI_Hs01_00115660) (siRNA pre-designed, Merck), KHDRBS1/Sam68 (ON-TARGETplus siRNA Smartpool L-020019-00, Horizon), KHDRBS2/SLM1 (target sequence GUGCAUGCGUCGCGCCUUU, purchased as custom siRNA from Merck), DHX9 (target sequence, AAGAAGUGCAAGCGACUCUAG, purchased as custom siRNA from Merck). Non-targeting scrambled siRNAs were used as negative control. RNA and proteins were extracted 72h after transfection, unless stated otherwise in the text. HEK293T cells were transfected using Lipofectamine 2000 Transfection Reagent (Cat. No. 11668019, Invitrogen) according to manufacturer’s instructions.

### Western blot analysis and antibodies

Cells were lysed in RIPA buffer (1% NP-40, 0.1% SDS, 150 mM NaCl, 50 mM Tris-HCl, pH=7.5, and 0.5% sodium deoxycholate) supplemented with 1% protease inhibitor cocktail (Sigma Aldrich), 0.5mM Na_3_VO_4_,1mM DTT. Equal amounts of proteins were separated on SDS-PAGE gels, transferred to PVDF membranes (BioRad) and probed with the following antibodies: anti-HNRNPC (Cat. No. 91327S, Cell Signaling Technology), anti-SART3 (Cat. No. GTX107684, GeneTex), anti-KHDRBS1/Sam68 (Cat. No. A302-110A, Bethyl Laboratories), anti-HSP90 (Cat. No. sc-13119, Santa Cruz Biotechnology), anti-histone H3 (Cat. No. 17168-1-AP, Proteintech), anti-DHX9 (Cat. No. A300-855A, Bethyl Laboratories). Anti-KHDRBS2/SLM1 was gently provided by Prof. Peter Scheiffele (University of Basel, Switzerland). Detection was achieved by using anti-mouse HRP-conjugated (Cat. No. NA931, Amersham) and anti-rabbit HRP-conjugated (Cat. No. NA934, Amersham) and visualized by Clarity Western ECL Substrate (Cat. No. 1705061, Bio-Rad).

### Total RNA extraction and quantitative real-time PCR (qPCR) analysesqPCR) analyses

Total RNA was extracted using miRNeasy mini kit (Cat. No. 217004, QIAGEN) according to manufacturer’s instructions. Total RNA was retro-transcribed with random primers using M-MLV reverse transcriptase (Promega) according to the manufacturer’s instructions. For circRNA detection, total RNA was treated with 2U RNaseR (Cat. No. RNR07250, Biosearch Technologies™) per μg of RNA for 20 minutes at 37°C. qPCR was carried out using LightCycler 480 SYBR Green I Master and the LightCycler 480 System (Roche), according to manufacturer’s instructions. Divergent primer pairs encompassing the back-splicing junction were used to detect circRNAs. The complete list of all primers used in this study is provided in Supplementary Table S19.

### UV cross-Linking and Immunoprecipitation (CLIP) assay

CLIP assays were performed as described in (41). D341 cells were UV-irradiated on ice (100 mJ/cm²). Following irradiation, cells were pelleted by centrifugation at 4000 rpm for 5 minutes. The resulting pellet was lysed on ice for 10 minutes in lysis buffer containing 50mM Tris (pH 8.0), 100mM NaCl, 1% NP-40, 1mM MgClC, 0.1mM CaCl_2_, 0.5mM Na_3_VO_4_, 1mM DTT, a protease inhibitor cocktail (Sigma-Aldrich), and RNase inhibitor (Promega). Lysates were briefly sonicated and subsequently treated with RNase-free DNase (Ambion) for 3 minutes at 37°C. After DNase treatment, samples were centrifuged at 15.000 g for 3 minutes at 4°C. For input RNA extraction, 0.1 mg of total extract was incubated with Proteinase K at 55°C for 30 minutes, and RNA was purified using standard protocols. For immunoprecipitation, 1 mg of extract was diluted to 1 ml in lysis buffer and incubated with 3 μg of anti-hnRNP C antibody (Cat. No. 91327S, Cell Signaling Technology) or control IgGs, along with protein G magnetic Dynabeads (Life Technologies). RNase I (1000 IU, Ambion) was added, and samples were incubated for 2 hours at 4°C with continuous rotation. After stringent washes (41), 10% of the immunoprecipitate was retained as an IP control. The remaining fraction was digested with 50 μg of Proteinase K at 55°C for 1 hour, and RNA was isolated following standard procedures.

### Plasmid constructs

circSMN minigene was generated as previously described (7). For mutagenesis of hnRNP C binding site (BS) 1, a megaprimer containing 13T>A at BS1 was amplified using primers #1 and #2 and circSMN minigene as a template. Megaprimer and primers #3 were used to amplify 5’ end of circSMN minigenes containing the mutagenized BS1. For mutagenesis of hnRNP C BS2, a second megaprimer with 12T>A at BS2 was obtained with primers #3 e #4 and then used with primer #1 to amplify the 5’ end of circSMN with mutated BS2. Double mutant (BS1 and BS2) was generated by using the same megaprimer-based strategy and circSMN BS2 mutant as template. All PCR amplicons generated for the mutations were digested with KpnI and SalI restriction enzymes for cloning into initial circSMN plasmid digested with the same enzymes. PCR primers listed in Supplementary Table S19.

### Cellular fractionation experiments

Following siRNA transfection for 72h, HD-MB03 cell pellet collected from two 10 cm dishes was lysed in a buffer containing 10mM Tris-HCl (pH 7.5), 0.5% NP-40, and 150mM NaCl, supplemented with 2mM sodium orthovanadate, 1mM DTT, and 1% protease inhibitor cocktail (Sigma Aldrich) allowing plasma membrane disruption. After 15 minutes incubation on ice, the lysate was layered over a chilled sucrose cushion (24% w/v in lysis buffer, 2.5 volumes) and centrifuged at 14.000 rpm for 10 minutes at 4°C. The supernatant containing the cytoplasmic fraction was collected. 50µL of cytoplasmic fraction was treated with 5U RNaseIII (Cat. No. AM2290, Invitrogen) at 37°C for 30 minutes, while the same volume (50µL) was left untreated at 37°C for 30 minutes. Cytoplasmic fractions were then resuspended in 20 volumes of Qiazol (QIAGEN) and RNA extracted using miRNeasy mini kit (Cat. No. 217004, QIAGEN) according to manufacturer’s instructions. The nuclear pellet was gently rinsed with cold PBS, treated with RNase III for 30 minutes at 37°C and then resuspended in Qiazol for nuclear RNA extraction.

### Immunofluorescence assay

HD-MB03 cells were washed in PBS, fixed with 4% (v/v) formaldehyde for 15 minutes at room temperature, washed 3 times with PBS, permeabilised with 0.5% (v/v) Triton X-100 in PBS for 15 minutes at room temperature, and washed twice with PBS. Cells were incubated with blocking solution (3% BSA in PBS) for 45 minutes at room temperature before incubation with anti-dsRNA J2 antibody (Cat. No. 76651, Cell Signaling Technology) at 4°C overnight. Cells were washed 3 times with PBS before incubation with secondary antibody (Alexa Fluor 488, Cat. No. A11029, ThermoFisher) for 1h at room temperature. Cells were again washed 3 times with PBS and the nuclei stained using 4′,6′- diamidino-2-phenykindole (DAPI). Coverslips were mounted with Vectashield mounting medium. Images were analysed by ImageJ software.

### Cell viability assay

Cell viability was assessed by CellTiter 96® AQueous One Solution Cell Proliferation Assay ([3-(4,5-dimethylthiazol-2-yl)-5-(3-carboxymethoxyphenyl)-2-(4-sulfophenyl)-2H- tetrazolium, inner salt; MTS, Cat. No. G3582 Promega] and performed in 96-well plates. For D341 cell line, 10000 cells/well were seeded. For HD-MB03 cell line, 3000 cell/well were seeded. The assay was performed by adding 20 μl of the CellTiter 96® AQueous One Solution Reagent directly to culture wells, incubating for 2-4h and then recording absorbance at 490 nm with a 96-well plate reader (iMark™ Microplate Absorbance Reader, BioRad). The quantity of formazan product as measured by the amount of 490 nm absorbance was directly proportional to the number of living cells in culture.

### Spheroid assay

D341 cells were transfected with siHNRNPC#1, siHNRNPC#2, or control siRNA for 24 hours. Cells were then trypsinised, resuspended in MEM full media containing 0.24% methyl cellulose (Cat. No. M7027, Merck), and seeded in round bottom ultra-low attachment 96 well plate (Cat. No. 7007, Corning) (250 cells per well). Plates were centrifuged at 100g for 5 minutes to allow cells to settle at the bottom of the wells. Spheroids growth were monitored at different time points using microplate multichannel automated imaging Celigo Image Cytometer (Nexcelom Bioscience) to assess changes in area and fluorescence intensity. It is important to note that the average mean fluorescent intensity (MFI) represents the averages of the MFI of the four replicate spheroids.

### RNA-seq samples alignment and back-splicing events calling in Group 3 MB patients

Calling of BS events was done using two distinct methods: CIRCexplorer2 version 2.3.8 (24) and CIRI version 2.1.1 (25). For the CIRCexplorer2 pipeline, EGA FASTQ samples (EGAD00001004958 dataset; DACO-1766 approved application ID at International Cancer

Genome Consortium platform) from MB, FC, and AC were downloaded from https://ega-archive.org and initially aligned to detect chimeric junctions using the STAR algorithm version 2.7.9a (42), with the following settings:

STAR \

--readFilesIn fastq1 fastq2 \

--runThreadN 16 \

--genomeDir STAR_HG19 \

--readFilesCommand zcat \

--outSAMtype BAM SortedByCoordinate \

--quantMode GeneCounts \

--chimSegmentMin 10 \

--outFileNamePrefix EGAF0000xxxxxxx_xx

where “fastq1” and “fastq2” refer to the fastq.gz read pair files, “STAR_HG19” is a directory containing the FASTA chromosome files for the human genome assembly GRCh37 (hg19), and a custom prefix is used, in the form EGAF0000xxxxxxx_xx, reflecting the EGA sample identifier. BS events were directly called and annotated from STAR chimeric junctions using CIRCExplorer2:

CIRCexplorer2 parse -t STAR EGAF0000xxxxxxxChimeric.out.junction > EGAF0000xxxxxxxCE2_parse.log

CIRCexplorer2 annotate \

-r hg19_ref_all.txt \

-g GENCODEv19/GRCh37.p13.genome.fa \

-b back_spliced_junction.bed \

-o EGAF0000xxxxxxx_circularRNA.txt

The “hg19_ref_all.txt” file is a collection of transcripts loci from UCSC Known Genes, RefSeq, and Ensembl transcripts, retrieved from the UCSC Genome Browser (43) database (last access: June 2022). GENCODE data and GRCh37 assembly FASTA files were also retrieved from the UCSC Genome Browser database.

For the CIRI pipeline, GRCh37 genome FASTA was indexed, and raw reads were aligned, using the BWA-MEM algorithm (44). The whole CIRI pipeline, including annotation, was run using the following settings:

java -jar /apps/ciri/2.1.1/bin/CIRI_Full_v2.1.1.jar \

-1 fastq1.fastq.gz \

-2 fastq2.fastq.gz \

-r GENCODEv19/GRCh37.p13.genome.fa \

-a GENCODEv19/gencode.v19.annotation.gtf \

-t 16 \

-d EGAF0000xxxxxxx_xx \

-o EGAF0000xxxxxxx_xx

The overlap between CIRCexplorer2 and CIRI calls is our high confidence BS set.

### Back-splicing junction quantification and differential junction usage

For CIRIquant analysis, BWA, HISAT2, StringTie and Samtools were used, and CIRIquant was used for quantification of circRNAs with the default parameters.

For circRNA differential expression analysis between MB and FC or AC samples we used prep_CIRIquant to summarize the circRNA expression profile in all samples obtained from CIRIquant, then prepDE.py and CIRI_DE_replicate for circRNA differential expression analyses using default parameters.

### Linear splicing and back-splicing analyses in hnRNP C-depleted MB cells

For linear splicing analyses refers to (45,46).

Back-spliced junctions (BSJs) were identified using CIRI-full algorithm (47), CIRIquant (48) was used for quantification and differential analysis, with RNAse-treated and -untreated samples. Annotations of circRNAs was done using GRCh38 genome and GENCODE v32 annotations. circRNAs were detected in RNAse-treated samples, but quantification was done on untreated samples to evaluate and take into account the efficiency of RNAse treatment (CIRIquant option). A circRNA was considered expressed in one sample if it has at least 2 reads on BSJ. A circRNA was expressed in one condition if it is expressed in at least half of the samples of the condition. A unique circRNA is a circRNA expressed in one sample only. For circular to linear ratio, the value was calculated by CIRI, only circRNA detected in all samples were reported.

Introns flanking BSJs from upregulated and unregulated circRNAs were identified using FAST DB v2022_1 annotations.

*Alu* definition was downloaded from UCSC using the “rmsk” table from the hg38 database. Inter-intronic IR*Alu*s were defined as pair of *Alu* sequences with each *Alu* from a given pair on one the two BSJ flanking introns and on the opposite direction. Each *Alu* sequence was counted once (i.e., a given *Alu* was not shared in several inter-intronic IR*Alu*s).

The number of inter-intronic IR*Alu*s per intron between BSJ flanking introns was compared with those within pairs of introns with similar length from expressed genes and from genes with BSJ but not BSJ flanking introns (to avoid bias due to intron length variation).

Strong hnRNP C binding motif was defined as polyT with at least nine “T”. Intron sequences were screened for hnRNP C motifs using a Perl script. Distances between hnRNP C motifs and BSJ or *Alu* were determined using R and Perl scripts.

### Analysis of RNA binding motifs

Sequence analysis for enriched motifs in intronic regions flanking splice sites involved circRNA biogenesis in AC was performed using the sequence motif discovery algorithm STREME (26) (https://meme-suite.org/meme/). 4-7 nt long motifs statistically significantly enriched (p≤0.05) were searched by comparative analyses of 100 nt long sequences upstream of the 5’ and 3’ splice sites of BSJs with those of analogous sequences from introns involved in canonical splicing events. Comparison of enriched motifs with the database from Ray *et al*. (29) of known binding sites for RBPs was performed using the Tomtom tool (28).

### Statistical analyses

GraphPad Prism software was used for statistical analysis. Data are presented as means±SD unless stated otherwise. Comparisons between groups were made using Student’s t-test, Welch’s t-test, or ANOVA as appropriate and as indicated in figure legends.

## SUPPLEMENTARY INFORMATION ACKNOWLEDGEMENTS

We wish to thank Dr. Marika Guerra and Dr. Valerio Petrera for technical assistance, and Prof. Pamela Bielli and Dr. Cinzia Caggiano for critical reading of the manuscript and helpful suggestions. We also thank all members of the laboratory for fruitful discussion throughout the course of this study.

## AUTHORS’ CONTRIBUTIONS

V.P. and A.M. conceived the study. A.M., C.P., S.C. and M.G. performed the experimental work. M.P., C.N., N.R., P.D.L.G., F.P. and L.G. performed the computational analysis. A.M., C.P., S.C., G.T., F.C., F.N., C.S. and V.P. interpreted and discussed the data. A.M., C.S. and V.P. wrote the manuscript. All authors approved the final version of the manuscript.

## FUNDING

The research leading to these results has received funding from Associazione Italiana Ricerca sul Cancro, AIRC under MFAG 2020 - ID. 24767 project – P.I. Pagliarini Vittoria. This work was also supported by Ministry of University [P2022JLHZZ to V.P.], and Fondazione MiaNeri S.r.l. to V.P.. A.M. was supported by a post-doctoral fellowship from Fondazione Umberto Veronesi. C.P. was supported by post-doctoral fellowship from AIRC [Project code 28286]. Università Cattolica del Sacro Cuore contributed to the funding of this research project and its publication.

## AVAILABILITY OF DATA AND MATERIALS

All data generated or analyzed during this study are included in this article. RNA-seq data are available on GEO database (accession number: GSE304204).

## ETHICS APPROVAL AND CONSENT TO PARTICIPATE

Ethical approval was not applicable to this study, no patients were enrolled and no consent was required.

## CONSENT FOR PUBLICATION

All authors provided their consent to publish the study.

## COMPETING INTERESTS

The authors declare no competing interests.

## Supporting information

Supplementary Figures S1 to S5_Legends for Supplementary Tables S1 to S19

## Supplementary Materials

Supplementary Figs. S1 to S5

Legends for Supplementary Tables S1 to S19

## Other Supplementary Material for this manuscript includes the following

Supplementary Tables S1 to S19 (as separate single file)

## Notes

### Competing Interest Statement

The authors have declared no competing interest.

