## Supplementary Figures S1 to S5_Legends for Supplementary Tables S1 to S19 for "HnRNP C binding to inverted *Alu* elements protects the transcriptome from pre-mRNA circularization"

**Running title:** hnRNP C inhibits the biogenesis of circRNAs

**Keywords:** RNA-binding proteins, inverted repeated elements, circular RNA, back-splicing

**Corresponding Author:**

Vittoria Pagliarini, Department of Neuroscience, Section of Human Anatomy, Catholic University of the Sacred Heart, 00168, Rome, Italy;

[#]Current Address: *Integrated Omics Department, Novo Nordisk, 2860 Søborg, Denmark*

**The pdf file includes:**

**Figs. S1 to S5**

**Tables S1 to S19 (as separate single file)**

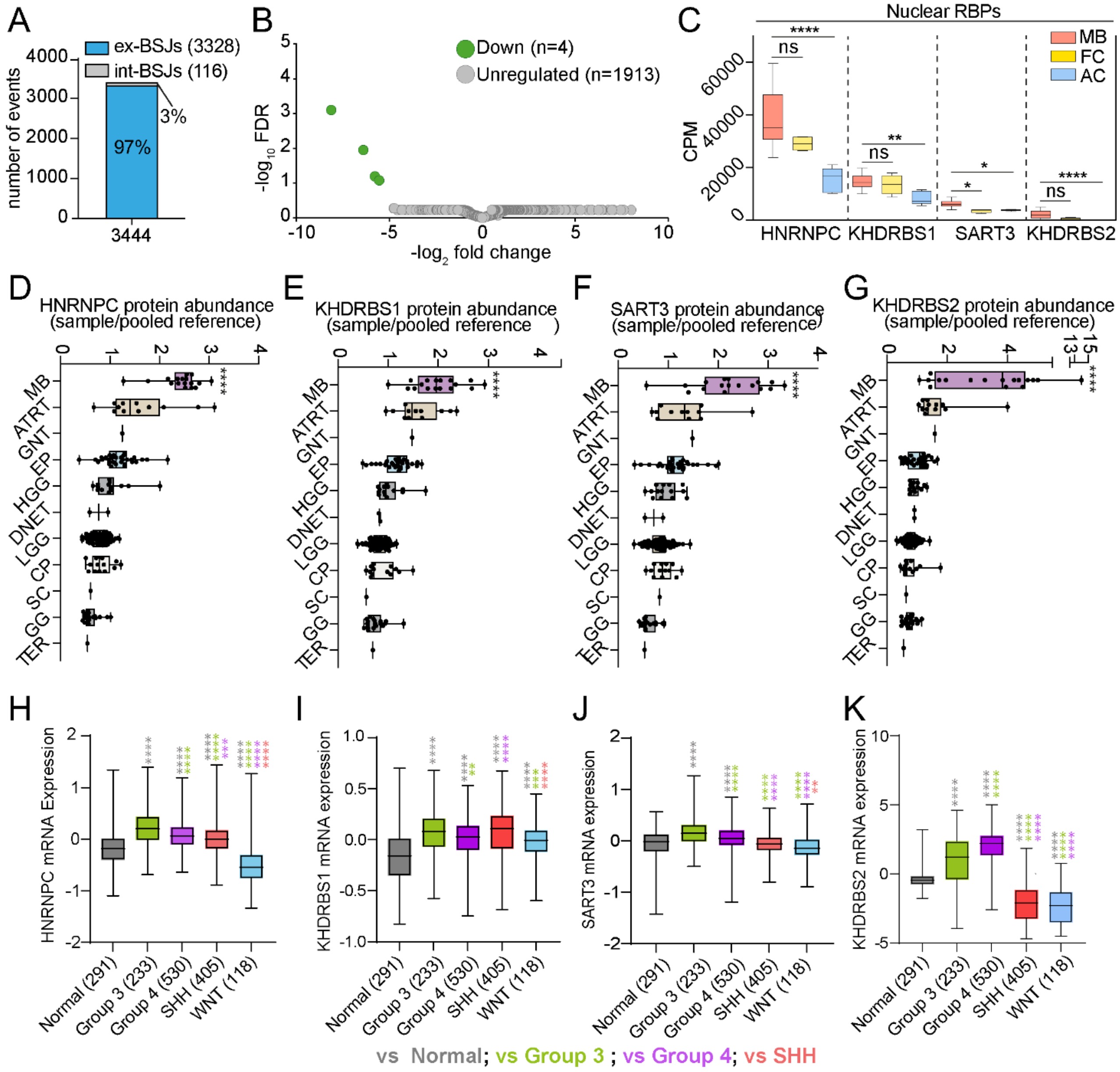

**Supplementary Fig. S1. CircRNA expression is extensively downregulated in Group 3 MB patients.** (**A**) Number of BSJs detected in Group 3 MB (MB), adult cerebellum (AC), and fetal cerebellum (FC) generated from exons or introns (exonic BSJs, ex-BSJs; intronic BSJs, int-BSJ). (**B**) Differential expression analysis of circRNAs detected in MB and FC. circRNAs differentially expressed (FDR<0,25) were highlighted in green. (**C**) mRNA expression levels (CPM, count per million reads) in EGAD00001004958 RNA-seq data of nuclear RBPs whose binding sites were enriched (Benjamini–Hochberg adjusted p-values; *p < 0.05, **p < 0.01, ****p < 0.0001). ns stands for not significant. (**D-G**) HNRNPC (**D**), KHDRBS1 (**E**), SART3 (**F**), and KHDRBS2 (**G**) protein abundance in Pediatric Brain Tumor Atlas database (CBTTC, Childhood Brain Tumor Tissue Consortium). Mass spectrometry data were retrieved from UCSC Xena platform (https://xenabrowser.net) (n=200). Medulloblastoma (MB, n=17), Atypical Teratoid Rhabdoid Tumor (ATRT, n=12), Glial-Neuronal Tumor (GNT, n=1), Ependymoma (EP, n=31), High-Grade Glioma/astrocytoma (WHO grade III/IV) (HGG, n=14), Dysembryoplastic Neuroepithelial Tumor (DNET, n=2), Low-Grade Glioma/astrocytoma (WHO grade I/II) (LGG, n=88), Craniopharyngioma (CP, n=13), Schwannoma (SC, n=1), Ganglioglioma (GG, n=19), Teratoma (TER, n=1) (One-way Anova; ****p-value summary < 0.0001). (**H-K**) HNRNPC (**H**), KHDRBS1 (**I**), SART3 (**J**), and KHDRBS2 (**K**) mRNA expression levels in Swartling MB (n=1286) and normal cerebellum (n=291) dataset. Data were retrieved from R2 platform (http://r2.amc.nl; Welch's t-test; **p < 0.01, ***p < 0.001, ****p < 0.0001). Only statistically significant differences are shown.

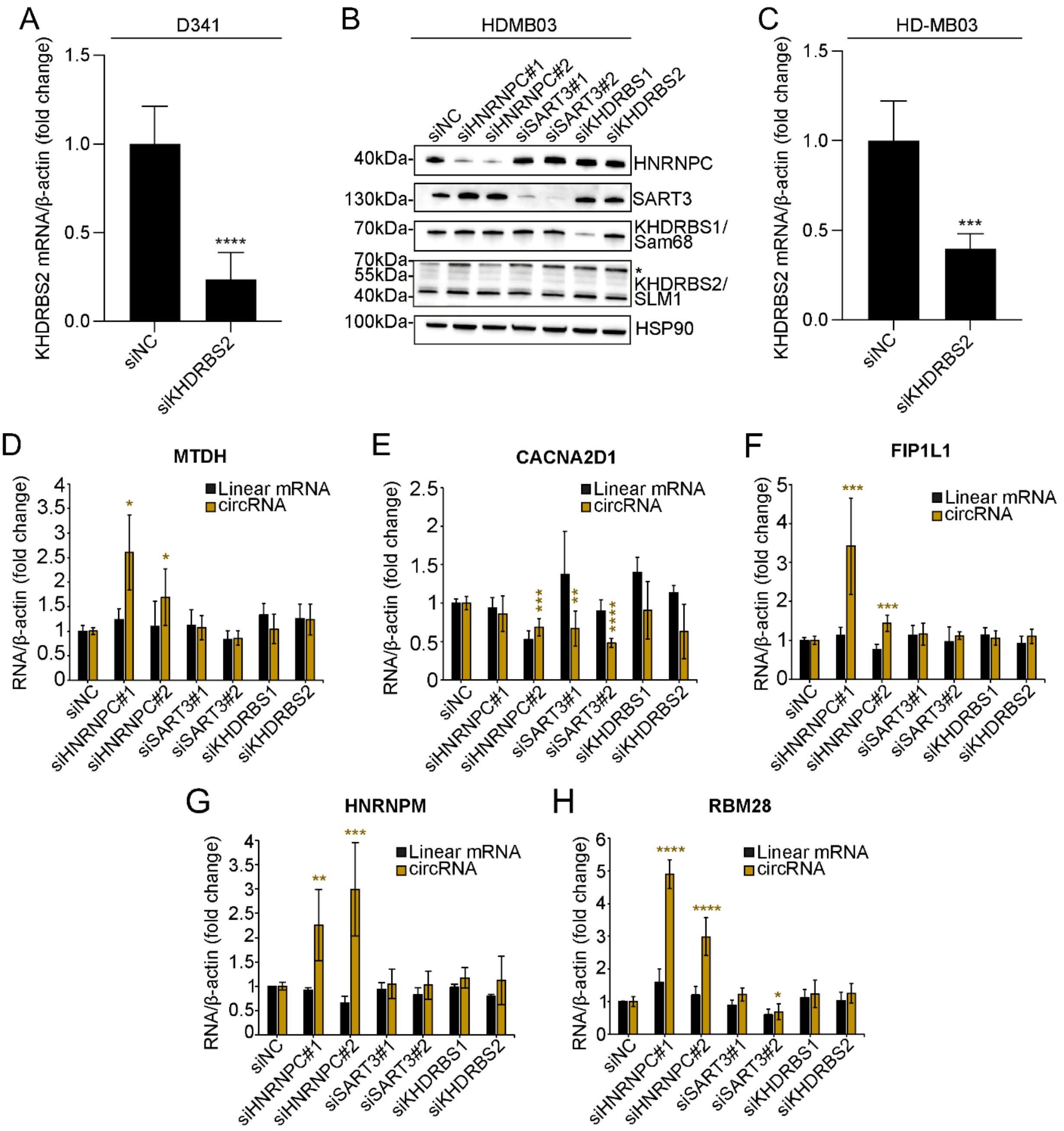


**Supplementary Fig. S2. HnRNP C represses the expression of circRNAs in Group 3 MB cells.** (**A**) qPCR analysis evaluating the expression of KHDRBS2 upon its silencing in D341 MB cells (n=3; mean with 95% CI; Student's t-test vs siNC; ****p < 0.0001). (**B**) Western blot analysis evaluating the expression of the selected RBPs upon their silencing in HD-MB03 MB cells. Asterisk (*) indicates a not-specific signal for KHDRBS1/Sam68 in KHDRBS2/SLM1 western blot. (**C**) qPCR analysis evaluating the expression of KHDRBS2 upon its silencing in HD-MB03 MB cells (n=3; mean with 95% CI; Student's t-test vs siNC; ***p < 0.001). (**D-H**) qPCRs detecting parental linear mRNAs and circRNAs having enrichment for HNRNPC, SART3, and KHDRBS1/2 binding sites in the flanking introns in HD-MB03 MB cells. #1 and #2 indicate two different siRNAs. siNC: siRNA negative control (n=3; mean ± SD; Student's t-test vs siNC; *p < 0.05, **p < 0.01, ***p < 0.001, ****p < 0.0001).

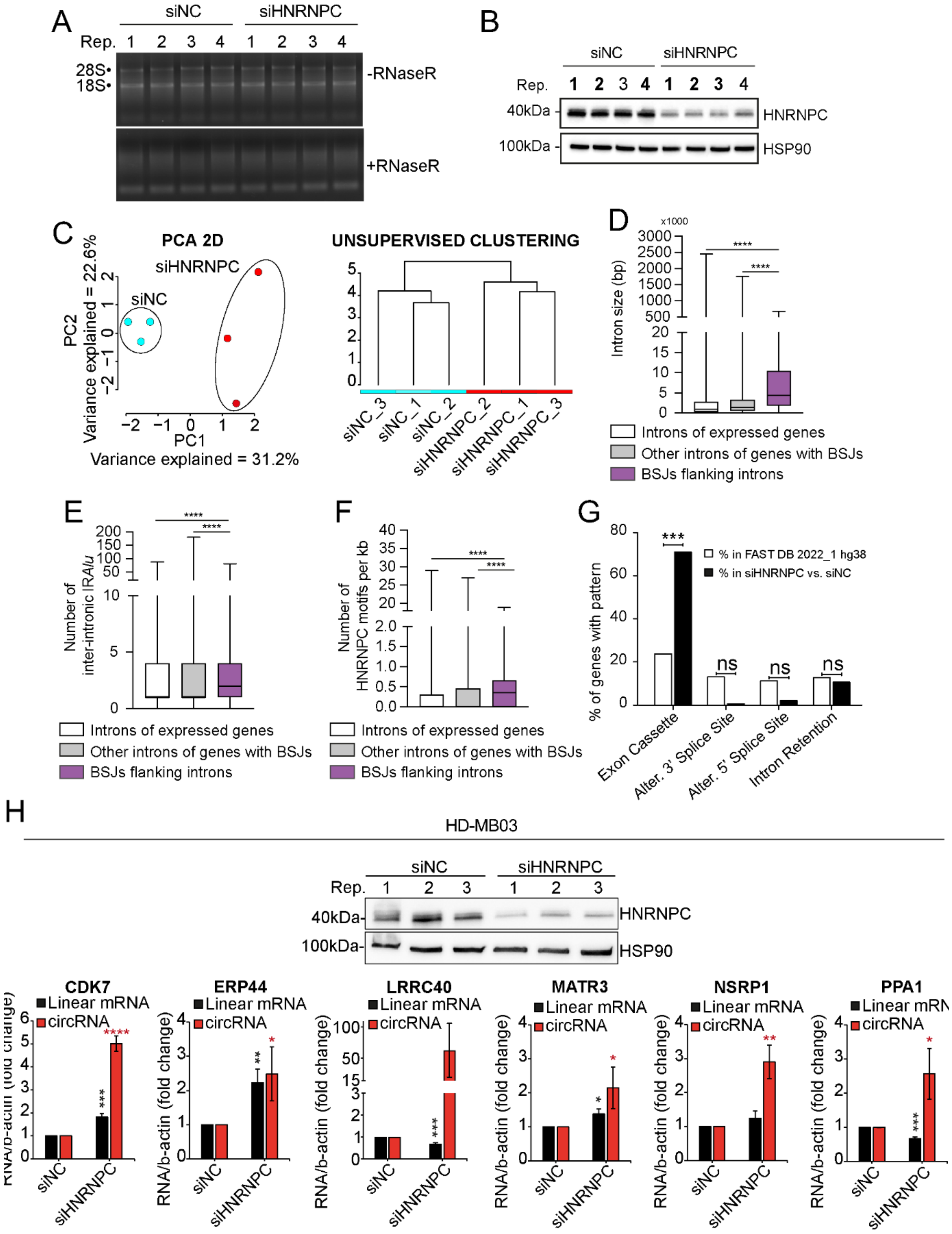
A
siNC
siHNRNPC
Rep. 1 2 3 4 1 2 3 4
28S
18S
-RNaseR
+RNaseR
B
siNC
siHNRNPC
Rep. 1 2 3 4 1 2 3 4
40kDa
HNRNPC
100kDa
HSP90
C
PCA 2D
siHNRNPC
siNC
PC2
Variance explained = 22.6%
PC1
Variance explained = 31.2%
UNSUPERVISED CLUSTERING
siNC_3
siNC_1
siNC_2
siHNRNPC_2
siHNRNPC_1
siHNRNPC_3
D
x1000
Intron size (bp)
****
Introns of expressed genes
Other introns of genes with BSJs
BSJs flanking introns
E
Number of inter-intronic IRAlu
****
Introns of expressed genes
Other introns of genes with BSJs
BSJs flanking introns
F
Number of HNRNPC motifs per kb
****
Introns of expressed genes
Other introns of genes with BSJs
BSJs flanking introns
G
% of genes with pattern
% in FAST DB 2022_1 hg38
% in siHNRNPC vs. siNC
***
ns
Exon Cassette
Alter. 3' Splice Site
Alter. 5' Splice Site
Intron Retention
H
HD-MB03
siNC
siHNRNPC
Rep. 1 2 3 1 2 3
40kDa
HNRNPC
100kDa
HSP90
CDK7
ERP44
LRRC40
MATR3
NSRP1
PPA1
Linear mRNA
circRNA
RNA/b-actin (fold change)
siNC
siHNRNPC

**Supplementary Fig. S3. HnRNP C is a general repressor of circRNA biogenesis in Group 3 MB cells.** (**A**) D341 MB cells were transfected with control (siNC) or HNRNPC (siHNRNPC) targeting siRNA for 48h. siHNRNPC#1 was used. (**B**) Western blot analysis for HNRNPC of samples in panel A. 3 out of 4 replicates (in bold) were used for RNAseq. (**C**) Principal component analysis (PCA; left panel) and unsupervised clustering (euclidean distance−method ward.D2; right panel) of RNAseq +RNaseR carried out in siRNA control (siNC) and siRNA HNRNPC (siHNRNPC) transfected D341 cells (3479 circRNAs). CIRI quant was used for circRNA detection and quantification. (**D-F**) Features of the flanking intron sequences of the detected circRNAs in control and HNRNPC-depleted D341 cells compared to either introns belonging to genes that do not generate circRNAs or introns that do not flank BSJs in genes that generate circRNAs (Student's t-test; ****$p < 0.0001$). (**G**) Percentages of events annotated in FAST-DB and of those regulated by HNRNPC depletion within each AS pattern (Student's t-test; ***$p < 0.001$). ns stands for not significant. (**H**) HD-MB03 MB cells were transfected with control (siNC) or HNRNPC (siHNRNPC) targeting siRNA for 72h. siHNRNPC#1 was used. Top panel: western blot showing silencing efficiency. Bottom panel: 6 up-regulated exonic circRNAs contained in the list of the top 8 most up-regulated circRNAs in HNRNPC-depleted D341 cells were validated in HD-MB03 cells (n=3; mean ± SD; Student's t-test vs siNC; *$p < 0.05$, **$p < 0.01$, ***$p < 0.001$, ****$p < 0.0001$).

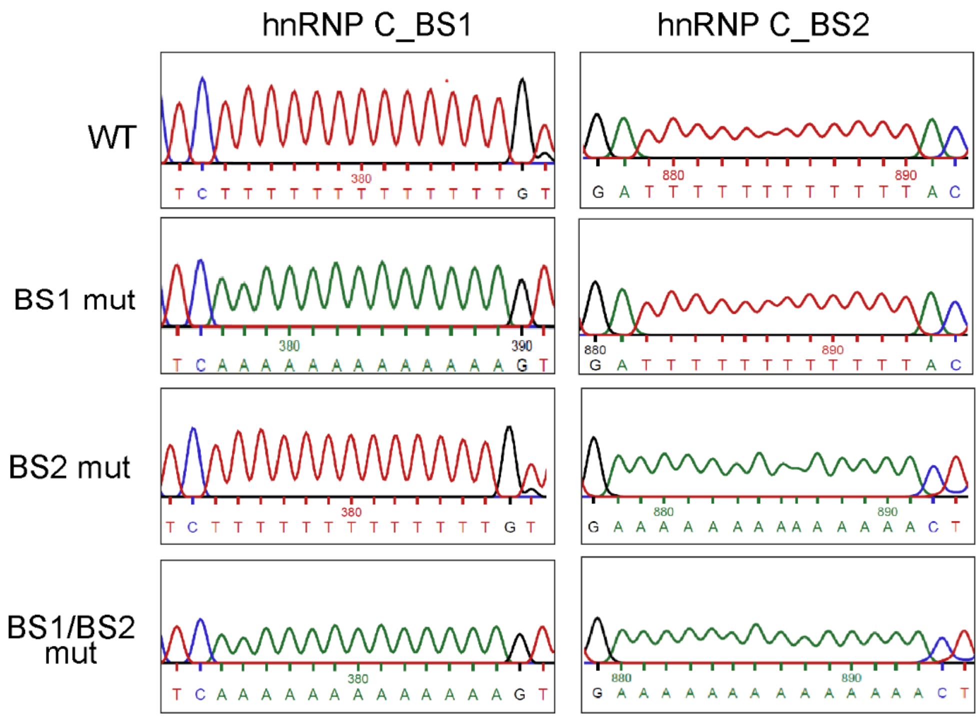


**Supplementary Fig. S4. Binding of hnRNP C is necessary to inhibit back-splicing events.** Sanger sequencing chromatograms of the wild-type (WT) and mutant circSMN9-6 minigenes. The WT trace confirms the native sequence of the HNRNPC binding sites, while chromatograms relative to HNRNPC-binding site (BS) 1, BS2, and BS1/BS2 mutants confirm that site-specific mutagenesis successfully occurred. In the mutants, all thymidines (T) within the HNRNPC binding motifs were replaced with adenines (A).

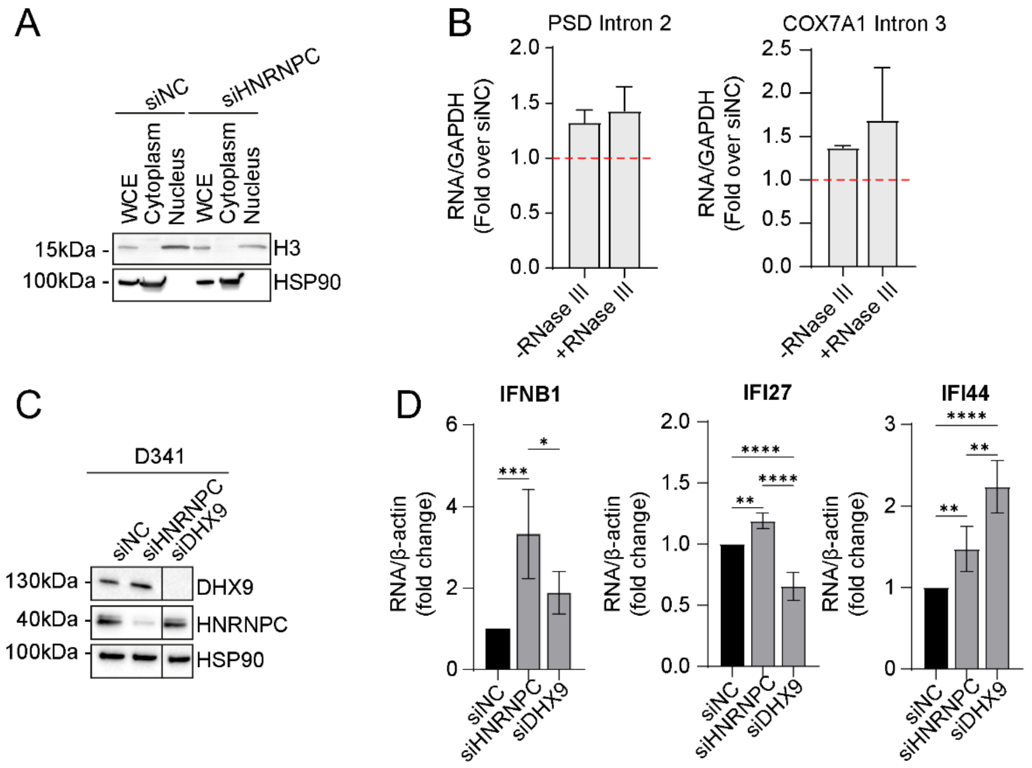


**Supplementary Fig. S5. HnRNP C represses the accumulation of intron-derived dsRNAs and spurious activation of innate immune response in Group 3 MB cells.** (**A**) Western blot analysis of subcellular fractionation performed in HD-MB03 MB cells treated with the indicated siRNAs. Histone H3 and HSP90 were used as markers for nuclear and cytoplasmic fractions, respectively**.** (**B**) qPCR analysis of introns not involved in back-splicing events (negative controls) are shown in the cytoplasmic fractions of hnRNP C-depleted HD-MB03 cells. Data are represented as fold change over siNC-treated cells set to 1, as indicated by the dashed red line. GAPDH was used as a loading control (n=3; mean ± SEM). (**C)** Western blot analysis of D341 cells treated with the indicated siRNAs. (**D**) qPCR analysis of immune-related genes in D341 cells treated with the indicated siRNAs (n=3/6; mean ± SD; One Way ANOVA, *$p < 0.05$, **$p < 0.01$, ***$p < 0.001$, ****$p < 0.0001$).

**Supplementary Table S1. Metadata and RNA-seq data files (EGAD00001004958 dataset) of subjects included in this study (healthy and Group 3 MB individuals) as reported in the International Cancer Genome Consortium (ICGC) Controlled Data database (https://ega-archive.org).**

**Supplementary Table S2. List of circRNAs detected in RNA-seq data from EGAD00001004958 dataset using CircExplorer2 and CIRI.**

**Supplementary Table S3. Differential expression analysis in Group 3 MB and AC of high confidence circRNAs detected in RNA-seq data from EGAD00001004958 dataset.**

**Supplementary Table S4. Differential expression analysis in Group 3 MB and FC of high confidence circRNAs detected in RNA-seq data from EGAD00001004958 dataset.**

**Supplementary Table S5. List of introns used for motif enrichment analysis (hg19).**

**Supplementary Table S6. Gene expression analyses of RNA-seq data from EGAD00001004958 dataset (Group 3 MB vs AC and Group 3 MB vs FC).**

**Supplementary Table S7. Genomic coordinates of EGAD00001004958-derived circRNAs tested in Group 3 MB cell lines.**

**Supplementary Table S8. All circRNAs detected in control siRNA and siHNRNPC#1 D341 cells.**

**Supplementary Table S9. Length and information of expressed genes' introns and circRNAs' flanking introns relative to Supplementary Figure S3D and Figure 4A (control siRNA and siHNRNPC#1 D341 cells).**

**Supplementary Table S10. Number of inter-intronic IR*Alu* in expressed genes' introns and circRNAs' flanking introns relative to Supplementary Figure S3E and Figure 4B (control siRNA and siHNRNPC#1 D341 cells).**

**Supplementary Table S11. Number of HNRNPC binding motifs in expressed genes' introns and circRNAs' flanking introns relative to Supplementary Figure S3F and Figure 4C (control siRNA and siHNRNPC#1 D341 cells).**

**Supplementary Table S12. siHNRNPC#1 vs control siRNA AS analysis (D341 cell line).**

**Supplementary Table S13. Differentially expressed genes in siHNRNPC#1 vs control siRNA gene expression analysis (D341 cell line).**

**Supplementary Table S14. Differentially expressed circRNAs in siHNRNPC#1 vs control siRNA (+RNaseR treated samples) expression analysis (D341 cell line).**

**Supplementary Table S15. Features (coordinates, length, *Alu*, IR*Alu*, HNRNPC binding motifs) of circRNAs flanking intron sequences (control siRNA and siHNRNPC#1 D341 cells).**

**Supplementary Table S16. Number of inter-intronic IR*Alu* (per kb) in circRNAs' flanking introns relative to Figure 4B (control siRNA and siHNRNPC#1 D341 cells).**

**Supplementary Table S17. Distance (bp) of HNRNPC binding motifs from *Alu* in circRNAs' flanking introns.**

**Supplementary Table S18. Distance (bp) of HNRNPC binding motifs from BSJs in circRNAs' flanking introns.**

**Supplementary Table S19. List of qPCR primers used in this study.**
